# Conformational Diversity and Substrate Specificity are Decoupled in Ancestral and Extant Glucokinases

**DOI:** 10.64898/2026.05.08.723840

**Authors:** Carolin Freye, Brian G. Miller

**Affiliations:** Department of Chemistry and Biochemistry, Laboratories of Molecular Recognition, Florida State University, 95 Chieftan Way, Tallahassee, FL 32306, USA

**Author notes:** **Corresponding Author** Brian G. Miller, 4005 Chemical Sciences Laboratory, Florida State University, Tallahassee, FL 32306-4390.

## Abstract

Multi-functionality in extant enzymes, including the ability to transform multiple substrates, is thought to arise, in part, from conformational flexibility. The hexokinase protein family represents a classic model system for investigating the relationship between substrate specificity and conformational change. Within this family, human glucokinase (hGCK) displays notable degrees of conformational heterogeneity, including an intrinsically disordered loop. The extent to which these structural features contribute to the breadth of hGCK’s substrate scope is unknown. Here, we investigate the substrate specificities of extant and ancestral glucokinases that span the evolutionary emergence of conformational heterogeneity in this family. We show that extant hGCK catalyzes the ATP-dependent phosphorylation of glucose, 2-deoxyglucose, mannose, glucosamine, fructose, allose and galactose with catalytic efficiencies ranging from 6.3 x 10^3^ M^-1^ sec^-1^ to 0.33 M^-1^sec^-1^. A glucokinase ancestor from early vertebrate evolution (vGCK), which also displays conformational heterogeneity and disorder, phosphorylates these same seven substrates with similar *k*_cat_/*K*_m_ values. An antecedent, chordate glucokinase (cGCK), which displays reduced conformational heterogeneity and lacks intrinsic disorder, also transforms these same substrates, but with higher overall catalytic efficiencies and markedly lower *K*_m_ values. Notably, however, the ratios of *k*_cat_/*K*_m_ values for individual substrate pairs, which define specificity, are unchanged for all three enzymes. Our results demonstrate that substrate specificity is not correlated with conformational diversity in GCKs and support a model in which the differences in catalytic efficiencies of various substrates arise from differences in the ability to form the ground state enzyme-carbohydrate binary complex.

---

In the late 1800s, Emil Fischer proposed the lock-and-key model of enzyme-substrate recognition, ascribing the origins of specificity to precise structural complementarity between the protein and the substrate.^1^ This model, which portrays the enzyme as a rather rigid entity, was later refined by Koshland when proposing the induced-fit model of protein-ligand interactions.^2^ The induced-fit model portrays the enzyme as more flexible and recognizes the role that ligand-induced protein structural rearrangements may play in substrate selection. Contemporary models view proteins in terms of freely equilibrating conformational ensembles, from which substrates can “select” a conformation for optimal enzymatic transformation.^3^ To varying degrees, the induced-fit and conformational ensemble models provide a framework for interpreting how conformational heterogeneity may contribute to the breadth of an enzyme’s specificity. Others, however, have argued that substrate specificity is independent of conformational changes for substrates that proceed through similar transition states and where similar rate-determining chemical or physical processes govern catalytic efficiency.^4^ Thus, the extent to which protein conformational diversity contributes to substrate specificity remains a matter of debate.

Some investigators have proposed that ancient enzymes were less specific than their extant counterparts.^5^ Indeed, an ancestral, “generalist” enzyme might offer the host an advantage by allowing a wider range of substrates to be processed using a smaller number of gene products. Consistent with this suggestion is the observation of phylogenetic homology between extant enzymes that operate on different substrates. These enzymes are postulated to have evolved through a process of gene duplication and sub/neofunctionalization, which could have originated with a broad specificity progenitor.^6^ However, many extant enzymes are known to function as generalists, retaining the ability to transform multiple substrates.^7^ Some have postulated that the conformational diversity and intrinsic disorder displayed by modern proteins could empower greater functional diversity.^8^ The extent to which ancestral proteins were less specific remains a topic of current discussion.

Structural studies of the hexokinase protein family provided some of the earliest evidence in support of the induced-fit theory of enzyme-substrate recognition.^2^ Hexokinases catalyze the ATP-dependent phosphorylation of various carbohydrates, undergoing an open-to-closed conformational transition upon substrate binding that prevents undesired hydrolysis of ATP in the absence of a suitable acceptor. Among the hexokinases, human hexokinase IV (also known as glucokinase, hGCK) displays a large degree of conformational heterogeneity.^9^ hGCK samples at least three major conformations — the super-open, open, and closed states (Figure 1).^10^ Unliganded hGCK predominantly exists in the super-open conformation, which is characterized by a wide opening angle between the large and small domains and a 30-residue intrinsically disordered loop.^9,11^ Upon binding, glucose shifts the conformational equilibrium to the open state.^11^ Association of the second substrate, MgATP^2-^, promotes formation of the closed, catalytically competent ternary complex.^11^ Like other sugar kinases, hGCK can transform multiple structurally related hexoses, but the extent to which conformational heterogeneity contributes to this broad substrate profile is unknown.^9^

**Figure 1.**
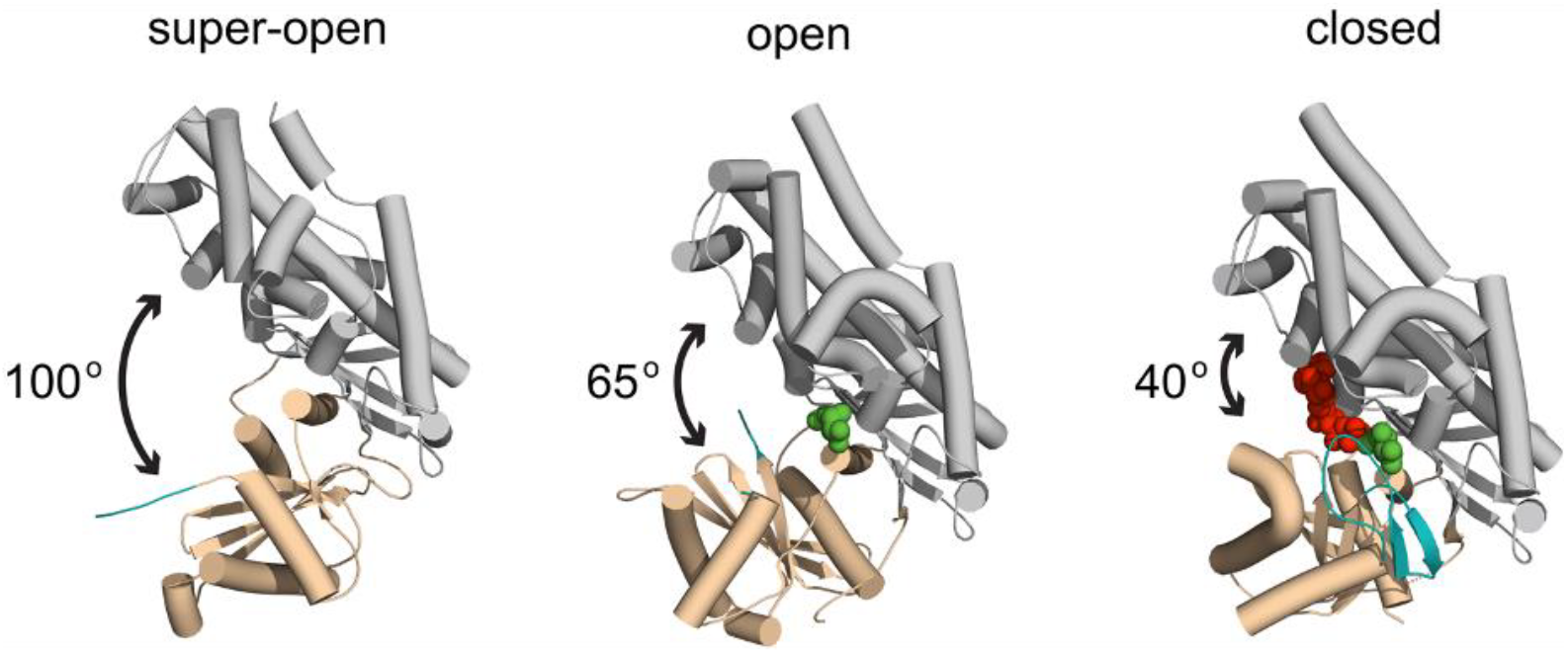
Conformations of human glucokinase. Unliganded hGCK predominantly adopts the super-open conformation with a large opening angle between the small (tan) and large (gray) domains and an invisible disordered loop (teal). Glucose (green) binding narrows the conformational ensemble promoting the open state. ATP (red) promotes the closed state and folding of the disordered loop in a β-hairpin (teal).

The ability of unliganded hGCK to sample multiple conformations endows this monomeric protein with a unique form of allostery, which manifests as a cooperative kinetic response to glucose concentrations.^9^ A recent phylogenetic study from our lab employed ancestral sequence reconstruction to investigate the molecular origins of kinetic cooperativity.^12^ The results revealed a discrete functional transition during early vertebrate evolution wherein an ancestral chordate GCK (cGCK) is non-cooperative and an ancestral vertebrate GCK (vGCK) displays a level of cooperativity equal to extant hGCK.^12^ The emergence of cooperativity correlates with a shift in several biochemical and biophysical traits that can be ascribed to an increase in conformational diversity. The transition from non-cooperative cGCK to cooperative vGCK coincided with an increased susceptibility to proteolysis within the disordered loop and increased backbone dynamics, particularly within the small domain, as revealed by hydrogen-deuterium exchange mass spectrometry.^12^ In addition, biomolecular NMR demonstrated that the unliganded, cooperative vGCK samples a broad conformational landscape, similar to hGCK, whereas the conformational ensemble of cGCK was narrower.^12^ Thus, these enzymes provide a useful system for investigating the potential links between substrate specificity and conformational diversity.

Here, we report the specificity profiles of hGCK, vGCK and cGCK to investigate whether the differences in conformational heterogeneity of these proteins correlate with the breadth of substrates accepted by each enzyme. We compare the substrate specificity of conformationally restricted, ancestral cGCK with the specificities of two conformational diverse glucokinase, vGCK and extant hGCK. We find that all three candidate enzymes transform the same set of carbohydrates, however, the non-cooperative cGCK does so with higher overall catalytic efficiencies for all substrates tested. We find that conformational heterogeneity coincides with higher *K*_m_/*K*_0.5_ values and lower overall catalytic efficiencies. Notably, the ratio of *k*_cat_/*K*_m_ values for any pair of substrates are largely unchanged for all enzymes, demonstrating that substrate specificity and conformational diversity are uncoupled in this protein family.

We investigated eight different hexoses as potential substrates for our extant and ancestral GCKs. Using an established enzyme-linked spectrophotometric assay, we attempted to determine *k*_cat_ and *K*_m_/*K*_0.5_ values for individual substrate using highly purified recombinant preparations of cGCK, vGCK, and hGCK. All three enzymes showed measurable activity toward seven common substrates: glucose, mannose, 2-deoxyglucose, glucosamine, fructose, allose and galactose (Table 1, Figures S1-32). We were only able to estimate *k*_cat_/*K*_m_ values for allose phosphorylation by vGCK and hGCK, and galactose phosphorylation by all three enzymes, using rate data from assays performed under sub-saturating concentrations (Figures S33-37). We observed apparent hexokinase activity with methyl-glucopyranose; however, analysis of the reaction products by mass spectrometry did not support the enzymatic production of methyl-glucopyranose 6-phosphate (Figures S38A). Instead, the results suggested that the observed activity originated from glucose present within the methyl-glucopyranose stock, which became observable due to the high carbohydrate concentrations used in our assays (Figure S38B). Thus, we excluded methyl-glucopyranose from our substrate profile. Similar mass spectrometry analysis confirmed the ATP-dependent synthesis of galactose 6-phosphate by vGCK, the lowest activity enzyme-substrate pair tested in this study (Figure S39).

**Table 1.** Kinetic parameters of various substrates transformed by cGCK, vGCK and hGCK.

|  | **cGCK** |  | **vGCK** |  | **hGCK** |  |
| --- | --- | --- | --- | --- | --- | --- |
| | $k_{\text{cat}}$ (s<sup>-1</sup>) | $K_m$ (mM) | $k_{\text{cat}}$ (s<sup>-1</sup>) | $K_{0.5}$ (mM) | $k_{\text{cat}}$ (s<sup>-1</sup>) | $K_{0.5}$ (mM) |
| **Glucose** | 45 ± 11 | 0.17 ± 0.03 | 41 ± 7 | 2.9 ± 0.4 | 56 ± 1 | 8.9 ± 0.5 |
| **Mannose** | 32.0 ± 0.1 | 0.19 ± 0.03 | 38 ± 7 | 11.2 ± 0.6 | 42 ± 14 | 15 ± 2 |
| **2-deoxyglucose** | 38 ± 2 | 0.29 ± 0.03 | 26 ± 3 | 38 ± 1 | 34 ± 1 | 72 ± 3 |
| **Glucosamine** | 10 ± 2 | 0.28 ± 0.04 | 9 ± 1 | 5 ± 1 | 5 ± 1 | 3.7 ± 0.4 |
| **Fructose** | 83 ± 7 | 3.1 ± 0.7 | 65 ± 2 | 210 ± 6 | 74 ± 28 | 322 ± 8 |
| **Allose** | 8 ± 1 | 75 ± 19 | >11<sup>a,b</sup> | 1167<sup>a,b</sup> | nd | nd |
| **Galactose** | >1.0 ± 0.2<sup>b</sup> | 1230 ± 25<sup>b</sup> | nd | nd | nd | nd |

Kinetic data were fit to the Michaelis-Menten or the Hill equation. Kinetic parameters are reported as the average ± standard deviation of two biological replicates with three technical replicates (nd: not determined).
<sup>a</sup> Kinetic parameters were estimated from one biological replicate.
<sup>b</sup> Kinetic parameters were estimated at sub-saturating carbohydrate concentrations (Figure S40).

Glucose was the preferred substrate for all three enzymes, as determined by the magnitude of its second-order rate constant for enzymatic transformation, *k*_cat_/*K*_m_ (Table 2). Substrate specificity is defined by the ratio of second-order rate constants for two competing substrates; a value we report here as the specificity coefficient. To quantify the specificities of cGCK, vGCK and hGCK, we calculated the specificity coefficient of each substrate compared to glucose (Table 2). This value represents the factor by which each enzyme prefers glucose over other hexoses. For all three enzymes, mannose was the second-best substrate based on the magnitude of *k*_cat_/*K*_m_ values, and this carbohydrate was characterized by the smallest specificity coefficient values (1.6 to 4). Similarly, for all three enzymes, galactose was the worst substrate, with *k*_cat_/*K*_m_ values < 1 M^-1^ sec^-1^. This carbohydrate produced the largest specificity coefficients, with values ranging from 19,000 to 300,000. Statistical analysis of all specificity coefficients of individual substrates across all three enzymes revealed that the only significant differences are found between cGCK and vGCK with 2-deoxyglucose and fructose (p < 0.05). The respective specificity coefficients between cGCK and hGCK are not significantly different for any substrate. Thus, these data demonstrate that the conformationally restricted cGCK ancestor and the conformationally diverse vGCK and hGCK enzymes share very similar specificity profiles.

**Table 2.** Catalytic efficiencies and specificity coefficients of substrates transformed by cGCK, vGCK and hGCK.

|  | cGCK |  | vGCK |  | hGCK |  |
| --- | --- | --- | --- | --- | --- | --- |
| | $k_{cat}/K_m$ ($M^{-1} s^{-1}$) | *Specificity coefficient* | $k_{cat}/K_{0.5}$ ($M^{-1} s^{-1}$) | *Specificity coefficient* | $k_{cat}/K_{0.5}$ ($M^{-1} s^{-1}$) | *Specificity coefficient* |
| **Glucose** | $2.7 (\pm 0.8) \times 10^5$ | - | $1.4 (\pm 0.3) \times 10^4$ | - | $6.3 (\pm 0.4) \times 10^3$ | - |
| **Mannose** | $1.7 (\pm 0.2) \times 10^5$ | $1.6 \pm 0.5$ | $3.4 (\pm 0.7) \times 10^3$ | $4 \pm 1$ | $3 (\pm 1) \times 10^3$ | $2 \pm 1$ |
| **2-Deoxyglucose** | $1.3 (\pm 0.1) \times 10^5$ | $2 \pm 1$ | $6.8 (\pm 0.9) \times 10^2$ | $21 \pm 5^*$ | $4.8 (\pm 0.2) \times 10^2$ | $13 \pm 5$ |
| **Glucosamine** | $3.6 (\pm 0.8) \times 10^4$ | $7 \pm 3$ | $2.0 (\pm 0.5) \times 10^3$ | $7 \pm 3$ | $1.3 (\pm 0.02) \times 10^3$ | $5 \pm 1$ |
| **Fructose** | $2.7 (\pm 0.7) \times 10^4$ | $10 \pm 4$ | $3.0 (\pm 0.1) \times 10^2$ | $47 \pm 11^*$ | $2.3 (\pm 0.9) \times 10^2$ | $27 \pm 12$ |
| **Allose** | $1.0 (\pm 0.3) \times 10^2$ | $3 (\pm 1) \times 10^3$ | $6.2 \pm 0.4^a$ | $2.3 (\pm 0.6) \times 10^3$ | $3.4 \pm 0.1^a$ | $2 (\pm 1) \times 10^3$ |
| **Galactose** | $0.8 \pm 0.2^a$ | $3 (\pm 1) \times 10^5$ | $0.23 \pm 0.04^a$ | $6 (\pm 1) \times 10^4$ | $0.33 \pm 0.02^a$ | $1.9 (\pm 0.1) \times 10^4$ |

Specificity coefficients marked with an asterisk displayed a statistically significant difference compared to cGCK ( $p < 0.05$ ).
<sup>a</sup> Catalytic efficiencies were estimated from the slope of enzyme activity versus sub-saturating carbohydrate concentration plots.

Despite little to no difference in substrate specificity, vGCK and hGCK display reduced catalytic efficiencies with all carbohydrates investigated compared to cGCK (Table 2). cGCK catalyzes glucose phosphorylation with an efficiency of 2.7 (± 0.8) x 10^5^ M^-1^sec^-1^, a value that is reduced by more than one order of magnitude for both vGCK and hGCK (1.4 ± 0.3 x 10^4^ M^-1^sec^-1^ and 6.3 ± 0.4 x 10^3^ M^-1^sec^-1^, respectively). Similarly, the catalytic efficiencies for mannose, 2-deoxyglucose, glucosamine, fructose and allose decreased for enzymes with increased conformational heterogeneity. To understand the origins of the reduced catalytic efficiencies observed for vGCK and hGCK, we examined the individual kinetic parameters for all substrates (Table 1). Enzymatic turnover, *k*_cat_, varies only slightly depending on the carbohydrate substrate under investigation. All three enzymes displayed lowest turnover numbers with glucosamine (5-10 sec^-1^), and highest *k*_cat_ values were observed with fructose (∼75 sec^-1^). The *k*_cat_ values for allose with vGCK and hGCK, and galactose for all candidate glucokinases, could not be determined because of an inability to reach saturating concentrations with these substrates. Overall, *k*_cat_ values remain mostly consistent across all candidate glucokinases regardless of conformational diversity (∼40 sec^-1^).

The increase in conformational heterogeneity present in vGCK and hGCK had pronounced effects on the midpoint of carbohydrate responsiveness, as reflected by the *K*_m_/*K*_0.5_ value. The *K*_m_ value for glucose increased from 170 ± 30 μM for cGCK to *K*_0.5_ values of 2.9 ± 0.4 mM and 8.9 ± 0.5 mM for vGCK and hGCK, respectively. Similar trends were observed for mannose, 2-deoxyglucose, and glucosamine. The *K*_m_ value of cGCK with fructose, 3.1 ± 0.7 mM, increased to 210 ± 6 mM for vGCK and 322 ± 8 mM for hGCK. By contrast, the differences in *K*_m_ values for ATP, determined under saturating concentrations of carbohydrates, were much smaller across all three enzymes (Table 3). This observation is consistent with the knowledge that substrate binding to hGCK, and other mammalian orthologs, is strictly ordered. Glucose binds first, inducing domain closure, followed by subsequent ATP association that involves smaller structural rearrangements.^9^

**Table 3.** *K*_m, ATP_ values of cGCK, vGCK, and hGCK with different carbohydrates.

| | cGCK $K_{m, \text{ATP}}$ (mM) | vGCK $K_{m, \text{ATP}}$ (mM) | hGCK $K_{m, \text{ATP}}$ (mM) |
| --- | --- | --- | --- |
| **Glucose** | $0.23 \pm 0.08$ | $0.41 \pm 0.07$ | $0.58 \pm 0.02$ |
| **Mannose** | $0.18 \pm 0.03$ | $0.69 \pm 0.05$ | $0.51 \pm 0.03$ |
| **2-deoxyglucose** | $0.21 \pm 0.07$ | $0.33 \pm 0.02$ | $0.46 \pm 0.01$ |
| **Glucosamine** | $0.93 \pm 0.02$ | $2.2 \pm 0.5$ | $2.7 \pm 0.6$ |
| **Fructose** | $0.30 \pm 0.04$ | $0.8 \pm 0.1$ | $0.58 \pm 0.06$ |
| **Allose** | $0.4 \pm 0.3$ | nd | nd |
| **Galactose** | $2.4 \pm 0.7^a$ | nd | nd |

Kinetic data were fit to the Michaelis-Menten equation. ATP $K_m$ values were reported from two biological replicates of individual preparation as the average ± standard deviation with three technical replicates each. nd: not determined, because we were not able to reach saturating carbohydrate conditions.
<sup>a</sup> Kinetic parameters were estimated at sub-saturating carbohydrate concentrations (Figure S41).

Together, our data demonstrate that the reduction in catalytic efficiency for all substrates observed between conformationally restricted and conformationally diverse enzymes is due to increases in carbohydrate *K*_m_/*K*_0.5_ values. This finding is consistent with the possibility that differences in catalytic efficiency arise from changes in microscopic rate constant(s) associated with the formation and decomposition of the enzyme-carbohydrate binary complex. These observations suggest a model in which a larger portion of the carbohydrate intrinsic binding energy must be used to shift the conformation of structurally heterogeneous, cooperative vGCK and hGCK during formation of the ground-state Michaelis-complex, thereby resulting in higher apparent *K*_m_ values for each substrate. This model comports with known, non-cooperative activating variants of hGCK, which narrow the conformational landscape by shifting the conformational ensemble toward the open state.^9^ These disease-associated activating hGCK variants are characterized by glucose *K*_m_ values that are 10 to 100-fold lower than those of the cooperative, wild-type enzyme.

Here, we probed the extent to which conformational diversity is correlated with substrate specificity in ancestral and extant GCKs. Intriguingly, all three enzymes investigated in this study catalyzed the phosphorylation of multiple substrates with strikingly similar specificity coefficients. Our data demonstrate that substrate specificity is decoupled from conformational heterogeneity in GCKs. Instead, conformational diversity and intrinsic disorder appear to tune the kinetic responsiveness of these enzymes to different ranges of carbohydrate concentrations, which likely coincides with differences in physiological levels. In future work, it will be interesting to compare the results reported here with substrate specificity profiles of other ancestral proteins that display changes in conformational diversity to test the generality of our findings.

## ASSOCIATED CONTENT

### Supporting Information

Protein sequences and steady-state kinetic data for hGCK, vGCK, and cGCK are provided as supporting information. In addition, the supporting information includes the Hill coefficients for hGCK and vGCK and spectra obtained via mass spectrometry. The following files are available free of charge. Supporting information.docx

## AUTHOR INFORMATION

### Author Contributions

The manuscript was written through contributions of all authors. All authors have given approval to the final version of the manuscript.

## TABLE OF CONTENTS GRAPHIC

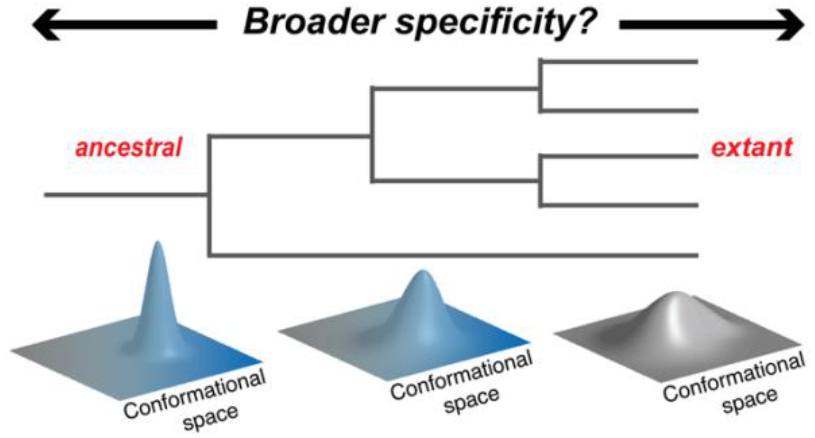

