## Supporting information for "Conformational Diversity and Substrate Specificity are Decoupled in Ancestral and Extant Glucokinases"

**TABLE OF CONTENT**

Materials and methods	1-2

Protein sequences	3

**Tables**

S1. Hill coefficients of vGCK and hGCK with various carbohydrate substrates	4

**Figures**

S1. Steady-state kinetic assays of cGCK with varying glucose concentrations fit to the Michaelis-Menten equation	5

S2. Steady-state kinetic assays of cGCK with varying ATP concentrations and saturating glucose concentrations fit to the Michaelis-Menten equation	6

S3. Steady-state kinetic assays of cGCK with varying mannose concentrations fit to the Michaelis-Menten equation	7

S4. Steady-state kinetic assays of cGCK with varying ATP concentrations and saturating mannose concentrations fit to the Michaelis-Menten equation	8

S5. Steady-state kinetic assays of cGCK with varying 2-deoxyglucose concentrations fit to the Michaelis-Menten equation	9

S6. Steady-state kinetic assays of cGCK with varying ATP concentrations and saturating 2-deoxyglucose concentrations fit to the Michaelis-Menten equation	10

S7. Steady-state kinetic assays of cGCK with varying glucosamine concentrations fit to the Michaelis-Menten equation	11

S8. Steady-state kinetic assays of cGCK with varying ATP concentrations and saturating glucosamine concentrations fit to the Michaelis-Menten equation	12

S9. Steady-state kinetic assays of cGCK with varying fructose concentrations fit to the Michaelis-Menten equation	13

S10. Steady-state kinetic assays of cGCK with varying ATP concentrations and saturating fructose concentrations fit to the Michaelis-Menten equation	14

S11. Steady-state kinetic assays of vGCK with varying glucose concentrations fit to the Hill equation	15

S12. Steady-state kinetic assays of vGCK with varying ATP concentrations and saturating glucose concentrations fit to the Michaelis-Menten equation	16

S13. Steady-state kinetic assays of vGCK with varying mannose concentrations fit to the Hill equation	17

S14. Steady-state kinetic assays of vGCK with varying ATP concentrations and saturating mannose concentrations fit to the Michaelis-Menten equation	18

S15. Steady-state kinetic assays of vGCK with varying 2-deoxyglucose concentrations fit to the Hill equation	19

S16. Steady-state kinetic assays of vGCK with varying ATP concentrations and saturating 2-deoxyglucose concentrations fit to the Michaelis-Menten equation	20

S17. Steady-state kinetic assays of vGCK with varying glucosamine concentrations fit to the Hill equation	21

S18. Steady-state kinetic assays of vGCK with varying ATP concentrations and saturating glucosamine concentrations fit to the Michaelis-Menten equation	22

S19. Steady-state kinetic assays of vGCK with varying fructose concentrations fit to the Hill equation	23

S20. Steady-state kinetic assays of vGCK with varying ATP concentrations and saturating fructose concentrations fit to the Michaelis-Menten equation	24

S21. Steady-state kinetic assays of hGCK with varying glucose concentrations fit to the Hill equation	25

S22. Steady-state kinetic assays of hGCK with varying ATP concentrations and saturating glucose concentrations fit to the Michaelis-Menten equation	26

S23. Steady-state kinetic assays of hGCK with varying mannose concentrations fit to the Hill equation	27

S24. Steady-state kinetic assays of hGCK with varying ATP concentrations and saturating mannose concentrations fit to the Michaelis-Menten equation	28

S25. Steady-state kinetic assays of hGCK with varying 2-deoxyglucose concentrations fit to the Hill equation	29

S26. Steady-state kinetic assays of hGCK with varying ATP concentrations and saturating 2-deoxyglucose concentrations fit to the Michaelis-Menten equation	30

S27. Steady-state kinetic assays of hGCK with varying glucosamine concentrations fit to the Hill equation	31

S28. Steady-state kinetic assays of hGCK with varying ATP concentrations and saturating glucosamine concentrations fit to the Michaelis-Menten equation	32

S29. Steady-state kinetic assays of hGCK with varying fructose concentrations fit to the Hill equation	33

S30. Steady-state kinetic assays of hGCK with varying ATP concentrations and saturating fructose concentrations fit to the Michaelis-Menten equation	34

S31. Steady-state kinetic assays of cGCK with varying allose concentrations fit to the Michaelis-Menten equation	35

S32. Steady-state kinetic assays of cGCK with varying ATP concentrations and saturating allose concentrations fit to the Michaelis-Menten equation	36

S33. Steady-state kinetic assays of cGCK at sub-saturating galactose concentrations	37

S34. Steady-state kinetic assays of vGCK at sub-saturating allose concentrations	38

S35. Steady-state kinetic assays of vGCK at sub-saturating galactose concentrations	39

S36. Steady-state kinetic assays of hGCK at sub-saturating allose concentrations	40

S37. Steady-state kinetic assays of hGCK at sub-saturating galactose concentrations	41

S38. Mass spectrometric analysis to probe the enzymatic production of methyl-glucoside 6-phosphate by cGCK	42

S39. Mass spectrometric analysis to probe the enzymatic production of galactose 6-phosphate by vGCK	43

S40. Steady-state kinetic assays of cGCK with varying galactose concentrations fit to the Michaelis-Menten equation	44

S41. Steady-state kinetic assays of cGCK with varying ATP concentrations and sub-saturating galactose concentrations fit to the Michaelis-Menten equation	45

**Materials and methods**

***Protein expression and purification.***

Protein sequences of cGCK and vGCK were inferred via ancestral sequence reconstruction as previously described.^1^ Wild-type hGCK and its ancestors, cGCK and vGCK, were expressed in *Escherichia coli* strain BL21(DE3) from pET-22b(+) vectors as N-terminal histidine tagged polypeptides (GenScript). Starter cultures were inoculated in Luria-Bertani media supplemented with ampicillin (150 µg/mL) at 37 °C overnight while shaking. Once the OD_600_ reached ~0.75, the temperature was reduced to 20 °C, and protein expression was induced with 0.75 mM isopropyl β-D-1-thiogalactopyranoside. Bacterial cultures were incubated at 20 °C overnight and harvested via centrifugation for 10 min at 6,000 x g and 4 °C.

Cell pellets were resuspended in chilled buffer A containing HEPES (50 mM, pH 7.4), KCl (50 mM), imidazole (45 mM), dithiothreitol (10 mM), and glycerol (25% v/v). Cells were lysed by French press at 1100 psi or by sonication. The crude cell lysates were clarified by centrifugation at 25,000 x g for 30 min at 4 °C and loaded onto a 5 mL HisTrap FF affinity column (Cytiva Life Sciences) pre-equilibrated with buffer A. The column was washed with 10 column volumes of buffer A to remove nonspecifically bound proteins. The recombinant glucokinases were eluted in buffer B containing HEPES (50 mM, pH 7.4), KCl (50 mM), imidazole (400 mM), dithiothreitol (10 mM), and glycerol (25% v/v). The eluate was dialyzed against potassium phosphate buffer (25 mM, pH 8.0) containing KCl (25 mM), and dithiothreitol (10 mM) overnight at 4 °C. Prior to size exclusion chromatography (SEC), the gel-filtration column (BioRad FPLC, Superdex 200 10/300 GL or ÄKTA FPLC, HiLoad 16/600 Superdex 200) was pre-equilibrated with SEC buffer containing HEPES (50 mM, pH 7.6) KCl (50 mM), dithiothreitol (10 mM). If necessary, protein samples were concentrated for injection using Amicon Ultra-15 Centrifugal Filter Units with a MWCO of 10 kDa (MilliporeSigma). Fractions displaying the highest levels of glucokinase activity were pooled.

***Enzyme assays.***

Glucokinase activity was assayed by coupling the production of ADP to the oxidation of NADH to NAD^+^ via the pyruvate kinase (5-20 units) - lactate dehydrogenase (10-15 units) coupled enzyme assay, and reaction progress was followed at 340 nm (Agilent Cary UV-Vis spectrophotometer). Lyophilized pyruvate kinase from rabbit muscle (MP Biomedicals, MilliporeSigma) was dissolved in chilled glycerol (25% v/v, pH 7.5). Lyophilized L-lactic dehydrogenase from rabbit muscle (MilliporeSigma) was dissolved in a sodium phosphate buffer (100 mM, pH 7.5) containing glycerol (25% v/v) and bovine serum albumin (1.0% w/v). Assays were performed in a Tris buffer (0.2 M, pH 7.6) containing NADH (0.5 mM), dithiothreitol (10 mM), phosphoenolpyruvate (5 mM), KCl (5 mM), carbohydrate (0.02–1500 mM), ATP (0.02–30 mM), and MgCl_2_ (2–31 mM). Carbohydrates were purchased from various sources (Fisher Scientific, MilliporeSigma, A2B Chem, and Calbiochem). Carbohydrate concentrations were varied while keeping the ATP concentrations at saturation to determine kinetic parameters, and vice versa. MgCl_2_ concentrations were kept at 1 mM above ATP concentrations. Assays were initiated via the addition of glucokinase. Data were fit to the standard Michaelis-Menten or the Hill equation by GraphPad Prism 6 depending on the enzyme and substrate under investigation. Hill coefficients are reported in Table S1. Kinetic parameters are reported as the average ± standard deviation of two independent protein preparations with three technical replicates performed for each data point (Table 1). The catalytic efficiencies reported in Table 2 were calculated by dividing the average *k*_cat_ values by the respective average *K*_m_ or *K*_0.5_ values listed in Table 1 unless otherwise stated. The standard deviations reported in Table 2 are the products of eq. 1 and the catalytic efficiency. The standard deviations reported for the specificity coefficients were calculated using the same approach. Statistical significance was determined using the student’s t-test.

σ= $\sqrt{{(\frac{st dev kcat}{avg kcat})}^{2}+{(\frac{st dev Km}{avg Km})}^{2}}$	(eq. 1)

***Mass spectrometry.***

The enzymatic production of galactose 6-phosphate and methyl-glucopyranose 6-phosphate, the two carbohydrates transformed with the lowest overall efficiency, was investigated using mass spectrometry. Samples containing enzyme (1.2-1.8 µM), galactose (250 mM) or methyl-glucopyranose (1 M), ATP (5 mM) and MgCl_2_ (6 mM) were incubated at room temperature overnight. Enzymes were removed from the reaction mixture via an Amicon Centrifugal Filter Unit with a MWCO of 10 kDa prior to sample analysis using an Agilent 6230 TOF-MS operating in negative ion mode.

**Protein sequences**

*Chordate GCK (cGCK)*

MHHHHHHGSGSMALREEKVELILDEFHLDNEELNEIMGRMHKEMEKGLRKETNEDATVKMLPTYVRSLPDGTESGDFLALDLGGTNFRVLLVKIKEGEELEGERKVEMKSQIYRIPEDVMTGTGEQLFDYIAECMADFLEKLGMKDRKLPLGFTFSFPCKQDGLDSASLITWTKGFSATGVEGKDVVKLLRDAIKRRGDFDMDIVAVVNDTVGTMMSCAFEDHDCLIGLIVGTGSNACYMEKMENVELLEGDKGEPNQMCINMEWGAFGDDGALDDFRTEYDREVDENSLNPGQQLYEKMISGMYMGELVRLVLLKLTKEGLLFGGKTSEELKTPGTFQTKYVSQIESDVPGDMTATLNILASLGLRHATEVDCEIVRQVCRAVSTRAAHLCAAGIAAVVNKMRRNRITVGVDGSVYKYHPTFKELMSETVDELTPGCDVKFMLSEDGSGKGAALITAVACRLAGK

*Vertebrate GCK (vGCK)*

MHHHHHHGSGSMLGRRSRMEGRKGEKVEQILSEFRLDKEELEEVMRRMQREMERGLRLETHEEASVKMLPTYVRSTPDGSEVGDFLALDLGGTNFRVMLVKVGEDEEGEWKVETKNQMYCIPEDVMTGTAEMLFDYIAECIADFLDKLNMKHKKLPLGFTFSFPVKHEDLDKGILINWTKGFTATGAEGNNVVELLRDAIKRRGDFDMDVVAMVNDTVATMISCYYEDHNCEIGMIVGTGCNACYMEEMRNVELVEGEEGRMCINMEWGAFGDSGELEEFRLEYDRKVDETSLNPGQQLYEKIISGKYMGELVRLVLLKLTNEGLLFGGKASEKLKTRGSFETKYVSQIESDDSGDMKQTYNILTTLGLQHPTELDCEIVRRVCQAVSTRAAHLCAAGMAAVVNKMRENRSQETLKITVGVDGSVYKLHPSFKDKFHAMVRELTPRCDITFIQSEEGSGRGAALISAVACKMACMGQ

*Human GCK (hGCK)*

MHHHHHHMVDDRARMEAAKKEKVEQILAEFQLQEEDLKKVMRRMQKEMDRGLRLETHEEASVKMLPTYVRSTPEGSEVGDFLSLDLGGTNFRVMLVKVGEGEEGQWSVKTKHQMYSIPEDAMTGTAEMLFDYISECISDFLDKHQMKHKKLPLGFTFSFPVRHEDIDKGILLNWTKGFKASGAEGNNVVGLLRDAIKRRGDFEMDVVAMVNDTVATMISCYYEDHQCEVGMIVGTGCNACYMEEMQNVELVEGDEGRMCVNTEWGAFGDSGELDEFLLEYDRLVDESSANPGQQLYEKLIGGKYMGELVRLVLLRLVDENLLFHGEASEQLRTRGAFETRFVSQVESDTGDRKQIYNILSTLGLRPSTTDCDIVRRACESVSTRAAHMCSAGLAGVINRMRESRSEDVMRITVGVDGSVYKLHPSFKERFHASVRRLTPSCEITFIESEEGSGRGAALVSAVACKKACMLGQ

**Tables**

**Table S1.** Hill coefficients of vGCK and hGCK with various carbohydrate substrates.

|  | **vGCK** | **hGCK** |
| --- | --- | --- |
| **Glucose** | 1.7 ± 0.1 | 1.76 ± 0.02 |
| **Mannose** | 1.73 ± 0.01 | 1.6 ± 0.1 |
| **2-deoxyglucose** | 1.20 ± 0.05 | 1.20 ± 0.05 |
| **Glucosamine** | 1.3 ± 0.1 | 1.47 ± 0.05 |
| **Fructose** | 1.6 ± 0.1 | 1.8 ± 0.2 |
| **Allose** | nd | nd |
| **Galactose** | nd | nd |

Kinetic data were fit to the Hill equation and Hill coefficients are reported as the average ± standard deviation of two bio-replicates with three technical replicates (nd: not determined).

**Figures.**

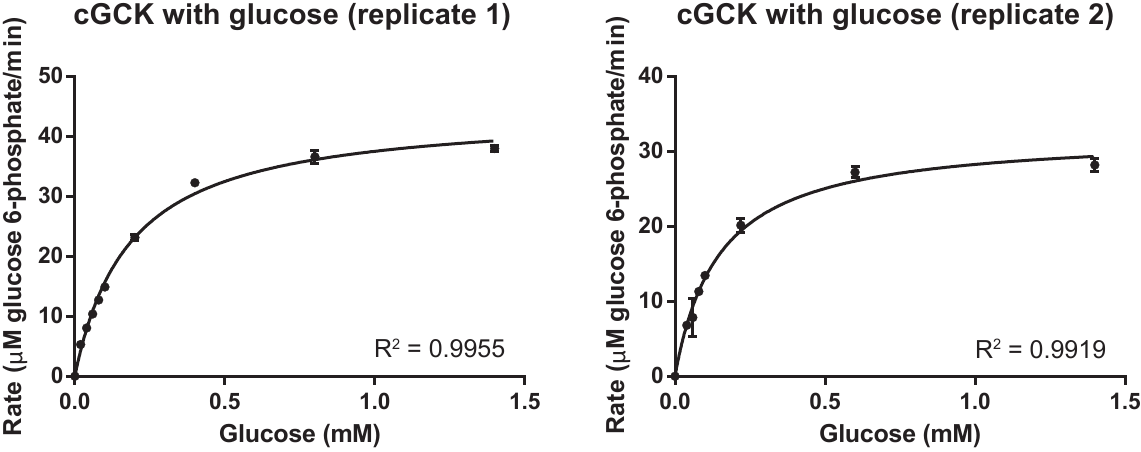

**Figure S1.** Two replicates of steady-state kinetic assays of cGCK with varying glucose concentrations at 10 mM ATP. Each data point represents the average ± standard deviation of three technical replicates. Data were fit to the Michaelis-Menten equation.

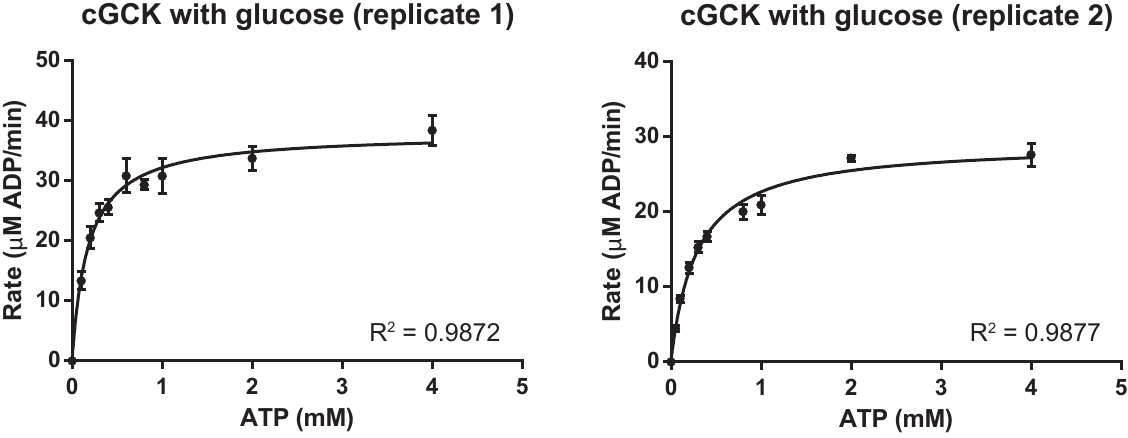

**Figure S2.** Two replicates of steady-state kinetic assays of cGCK with varying ATP concentrations at 2 mM glucose. Each data point represents the average ± standard deviation of three technical replicates. Data were fit to the Michaelis-Menten equation.

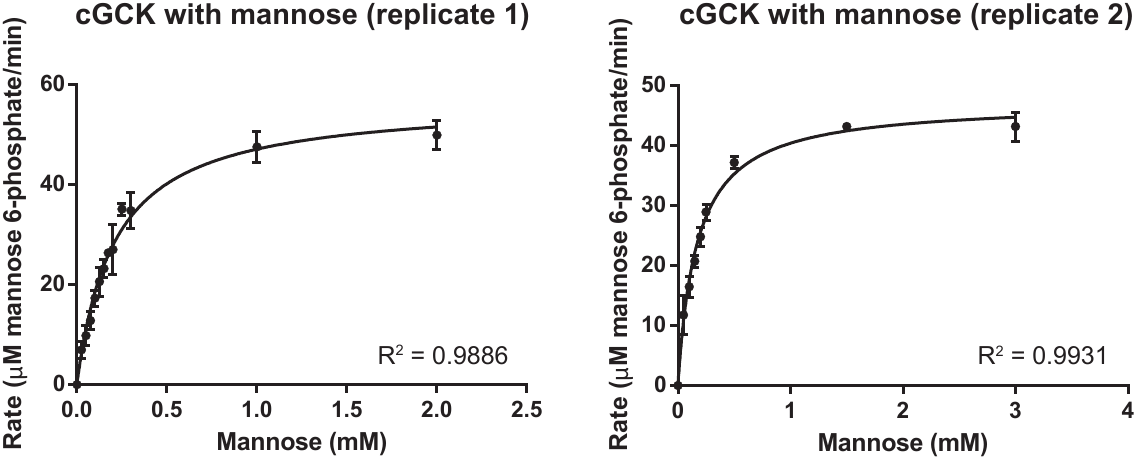

**Figure S3.** Two replicates of steady-state kinetic assays of cGCK with varying mannose concentrations at 10 mM ATP. Each data point represents the average ± standard deviation of three technical replicates. Data were fit to the Michaelis-Menten equation.

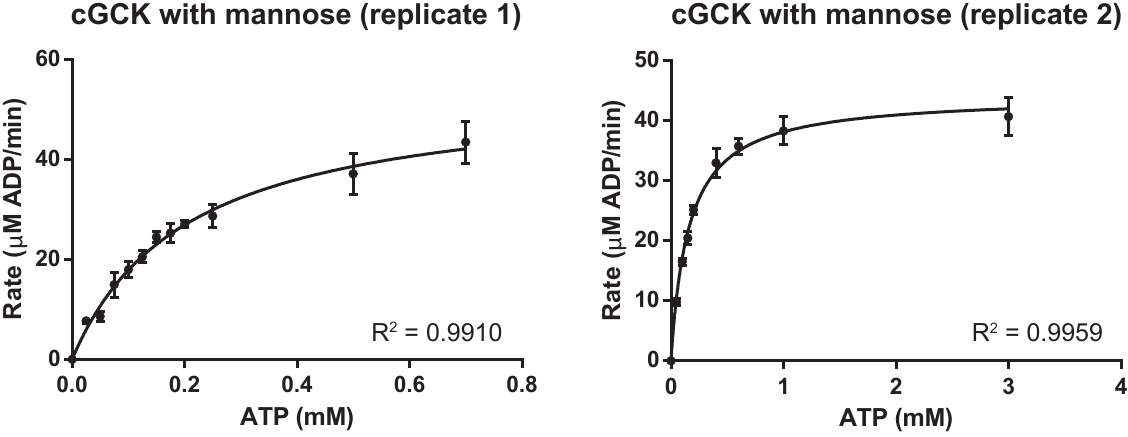

**Figure S4.** Two replicates of steady-state kinetic assays of cGCK with varying ATP concentrations at 5 mM mannose. Each data point represents the average ± standard deviation of three technical replicates. Data were fit to the Michaelis-Menten equation.

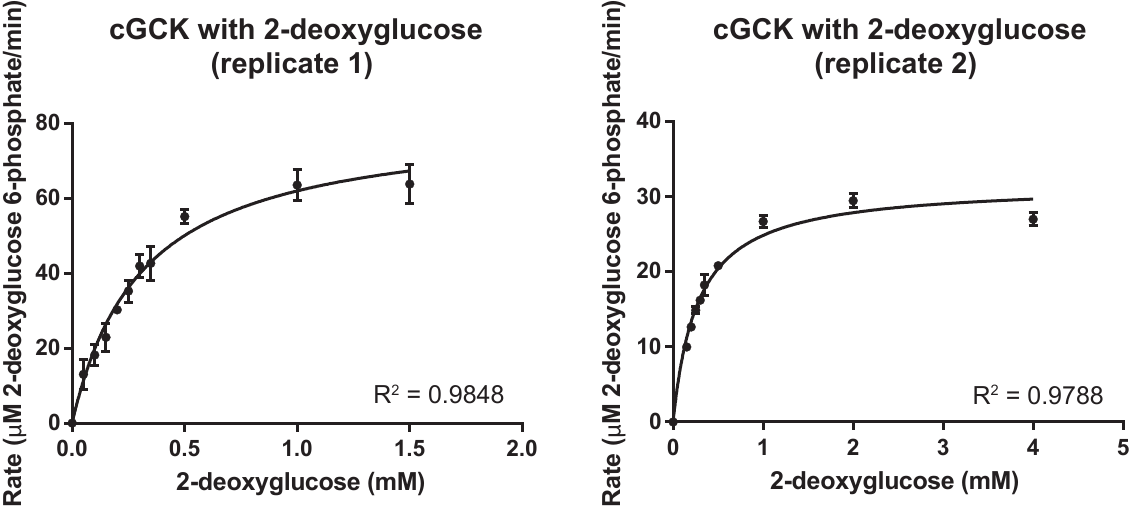

**Figure S5.** Two replicates of steady-state kinetic assays of cGCK with varying 2-deoxyglucose concentrations at 10 mM ATP. Each data point represents the average ± standard deviation of three technical replicates. Data were fit to the Michaelis-Menten equation.

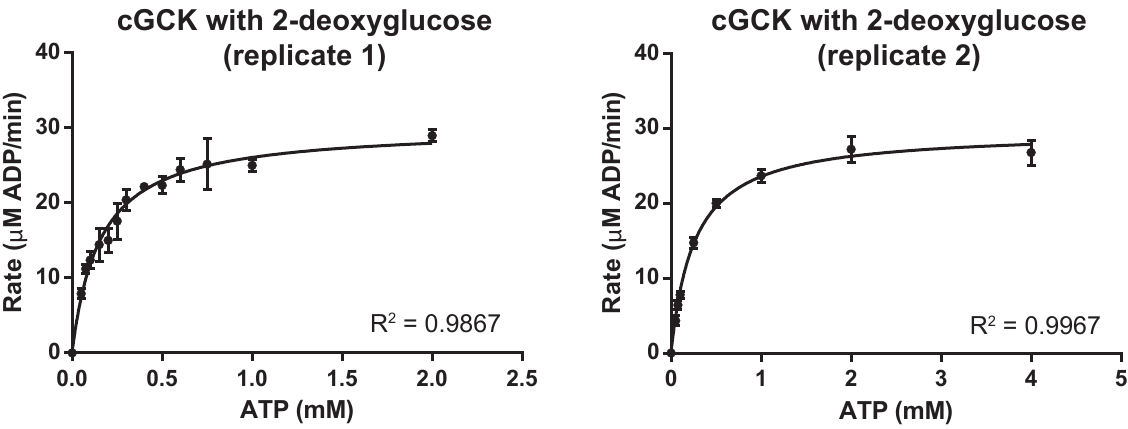

**Figure S6.** Two replicates of steady-state kinetic assays of cGCK with varying ATP concentrations at 20 mM and 10 mM 2-deoxyglucose for replicate 1 and 2, respectively. Each data point represents the average ± standard deviation of three technical replicates. Data were fit to the Michaelis-Menten equation.

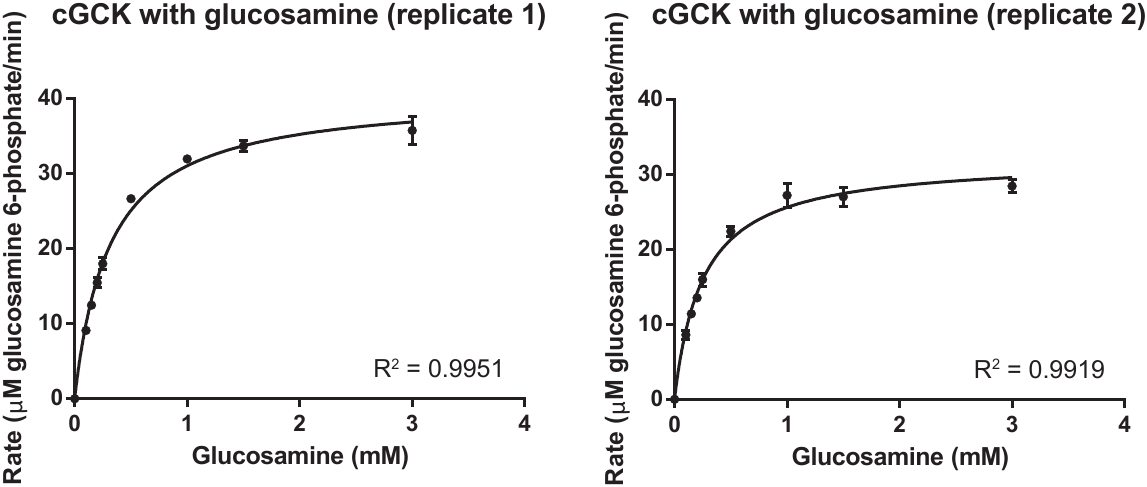

**Figure S7.** Two replicates of steady-state kinetic assays of cGCK with varying glucosamine concentrations at 10 mM ATP. Each data point represents the average ± standard deviation of three technical replicates. Data were fit to the Michaelis-Menten equation.

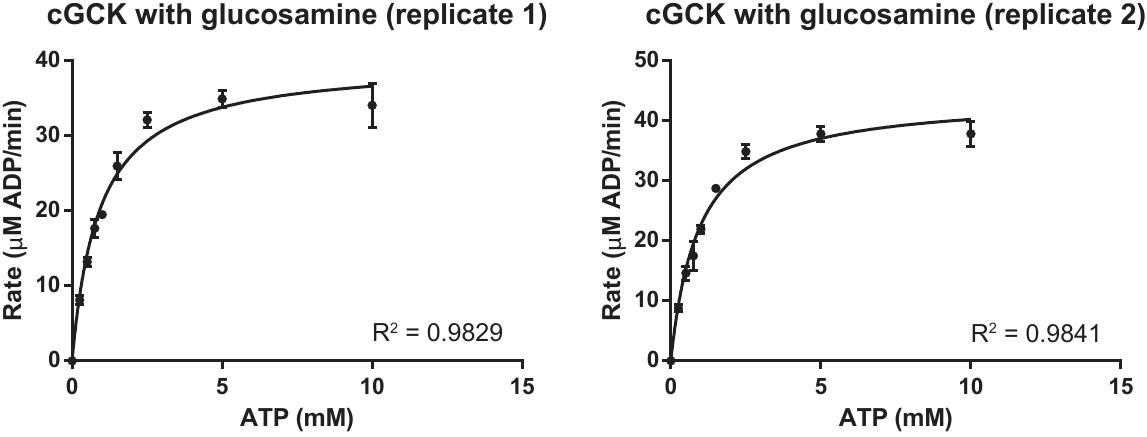

**Figure S8.** Two replicates of steady-state kinetic assays of cGCK with varying ATP concentrations at 6 mM glucosamine. Each data point represents the average ± standard deviation of three technical replicates. Data were fit to the Michaelis-Menten equation.

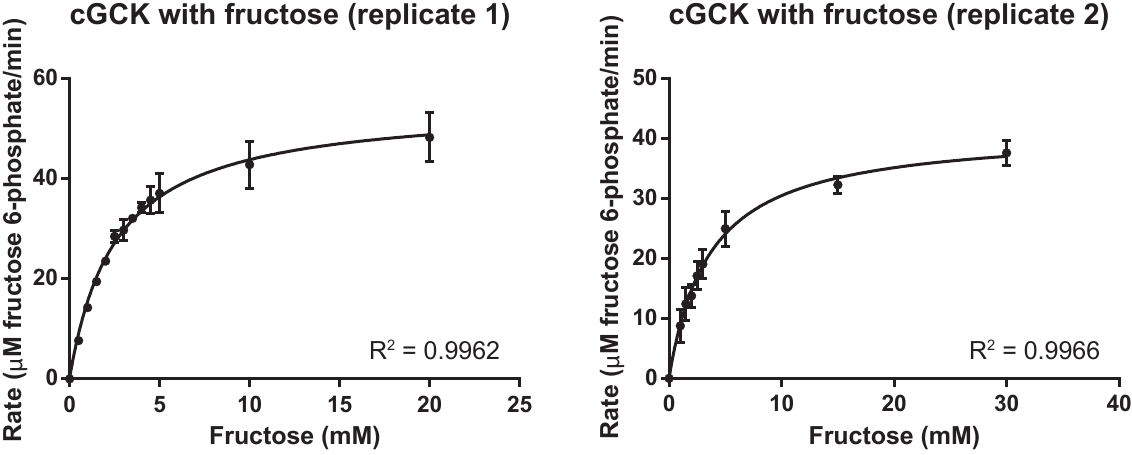

**Figure S9.** Two replicates of steady-state kinetic assays of cGCK with varying fructose concentrations at 10 mM ATP. Each data point represents the average ± standard deviation of three technical replicates. Data were fit to the Michaelis-Menten equation.

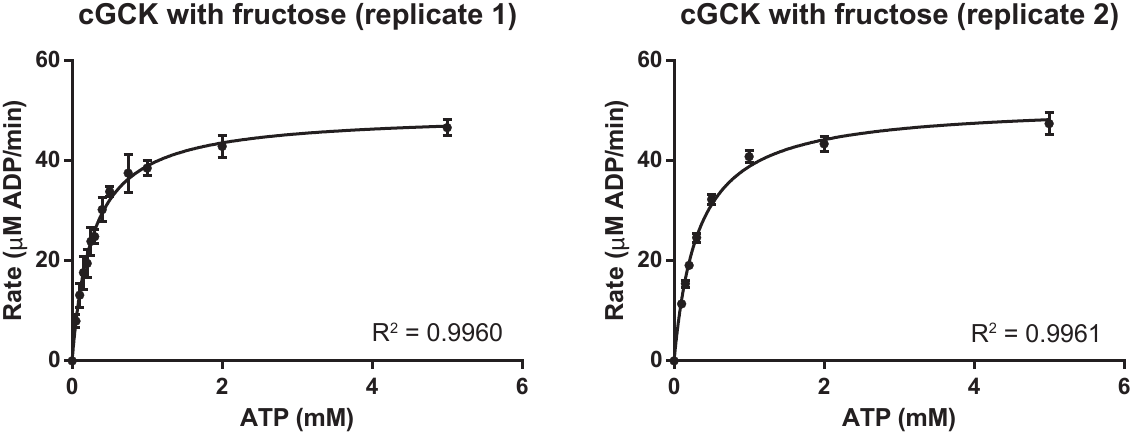

**Figure S10.** Two replicates of steady-state kinetic assays of cGCK with varying ATP concentrations at 50 mM fructose. Each data point represents the average ± standard deviation of three technical replicates. Data were fit to the Michaelis-Menten equation.

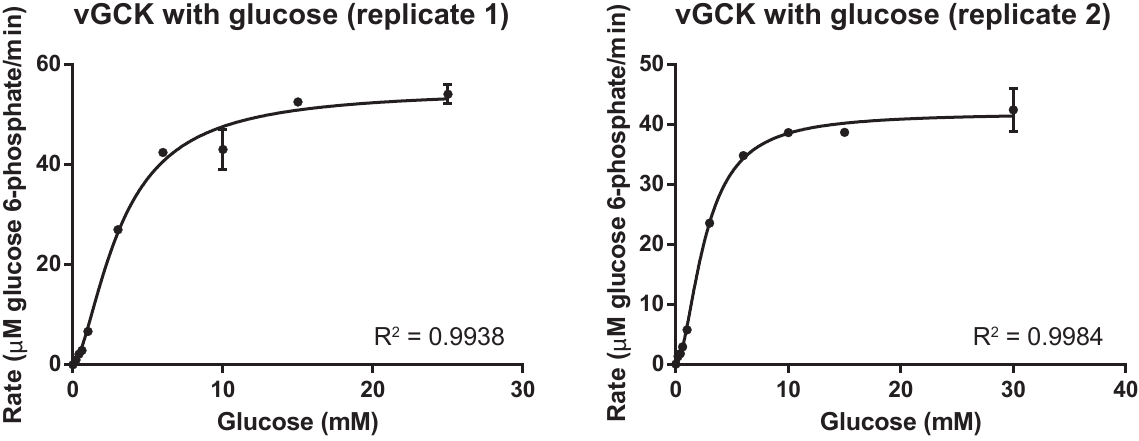

**Figure S11.** Two replicates of steady-state kinetic assays of vGCK with varying glucose concentrations at 10 mM ATP. Each data point represents the average ± standard deviation of three technical replicates. Data were fit to the Hill equation.

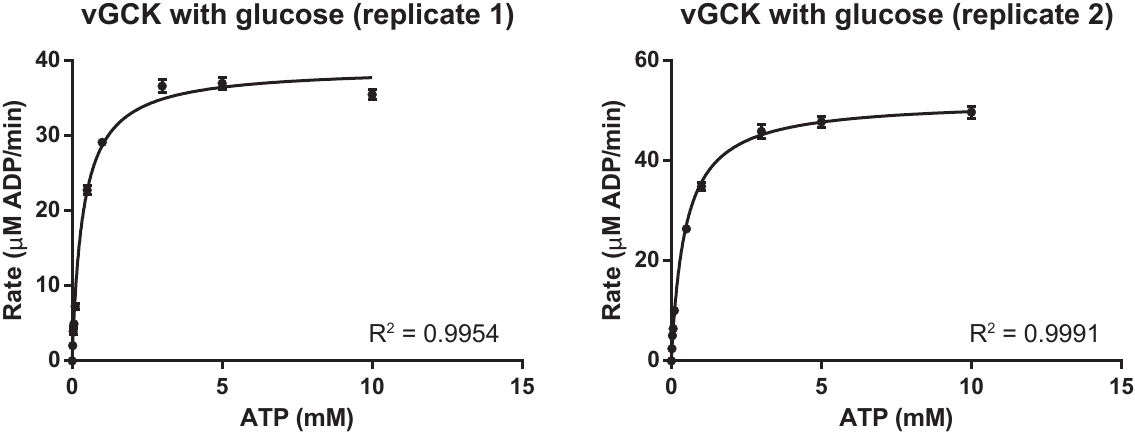

**Figure S12.** Two replicates of steady-state kinetic assays of vGCK with varying ATP concentrations at 100 mM glucose. Each data point represents the average ± standard deviation of three technical replicates. Data were fit to the Michaelis-Menten equation.

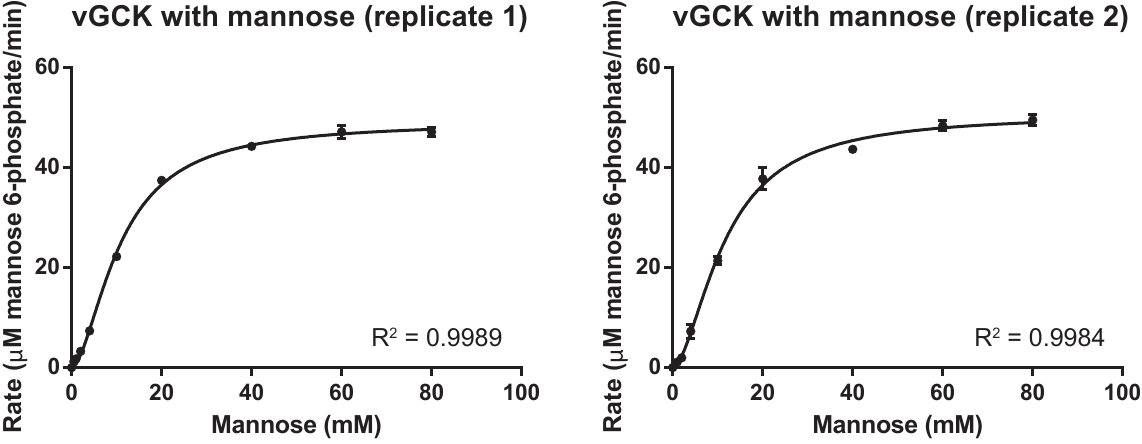

**Figure S13.** Two replicates of steady-state kinetic assays of vGCK with varying mannose concentrations at 10 mM ATP. Each data point represents the average ± standard deviation of three technical replicates. Data were fit to the Hill equation.

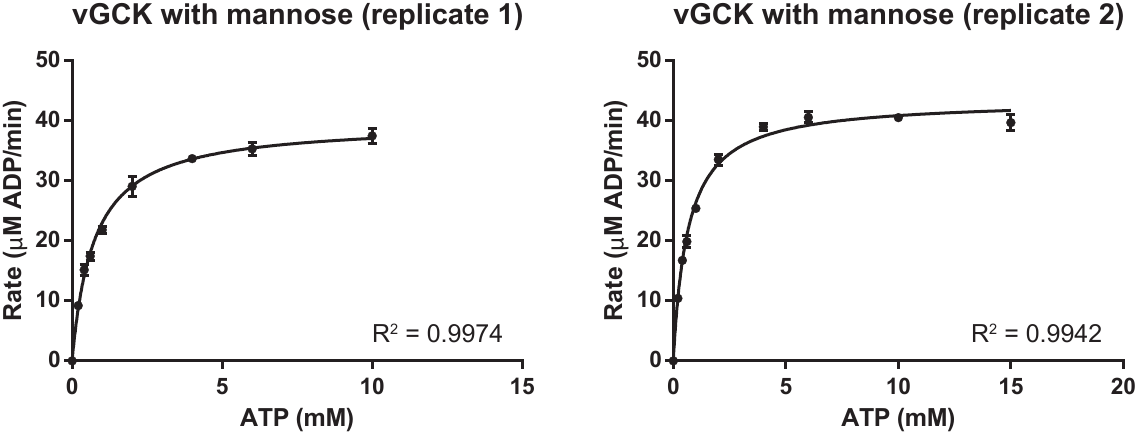

**Figure S14.** Two replicates of steady-state kinetic assays of vGCK with varying ATP concentrations at 100 mM mannose. Each data point represents the average ± standard deviation of three technical replicates. Data were fit to the Michaelis-Menten equation.

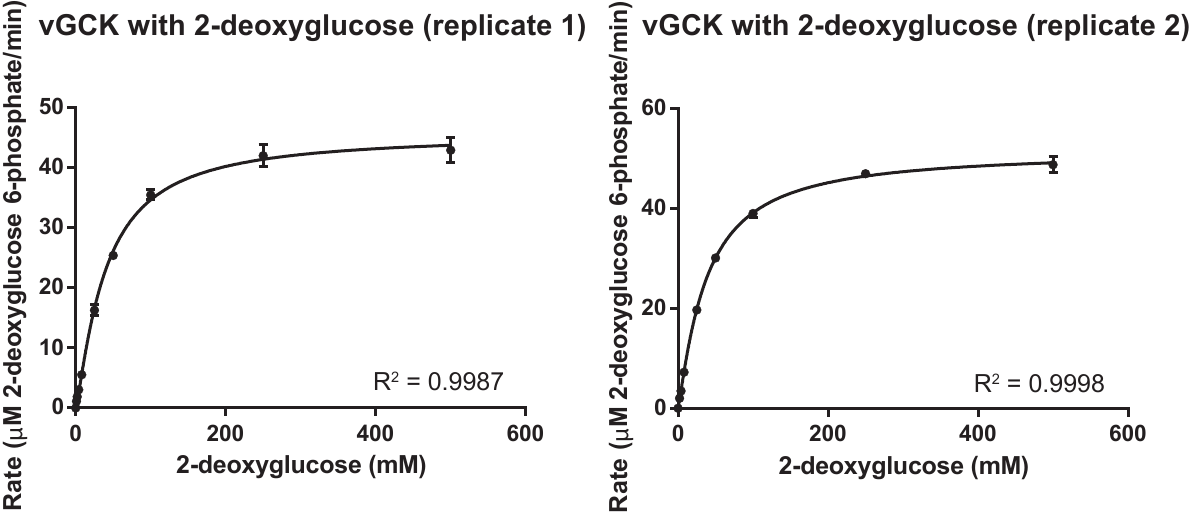

**Figure S15.** Two replicates of steady-state kinetic assays of vGCK with varying 2-deoxyglucose concentrations at 10 mM ATP. Each data point represents the average ± standard deviation of three technical replicates. Data were fit to the Hill equation.

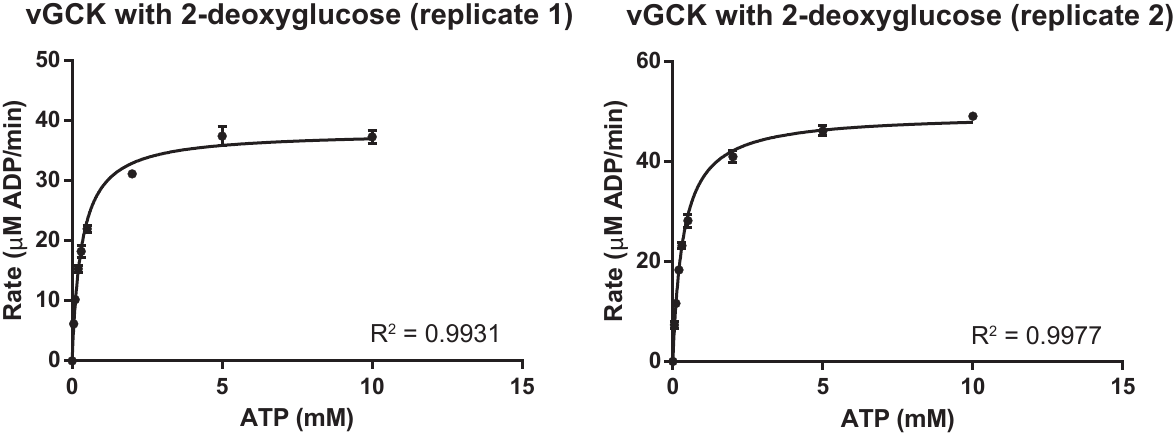

**Figure S16.** Two replicates of steady-state kinetic assays of vGCK with varying ATP concentrations at 500 mM 2-deoxyglucose. Each data point represents the average ± standard deviation of three technical replicates. Data were fit to the Michaelis-Menten equation.

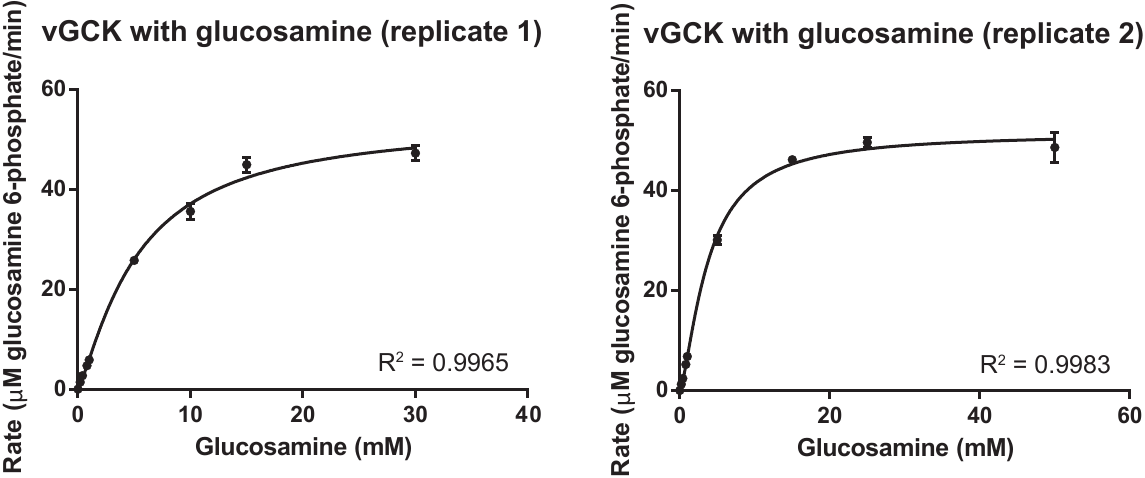

**Figure S17.** Two replicates of steady-state kinetic assays of vGCK with varying glucosamine concentrations at 10 mM and 20 mM ATP for replicate 1 and 2, respectively. Each data point represents the average ± standard deviation of three technical replicates. Data were fit to the Hill equation.

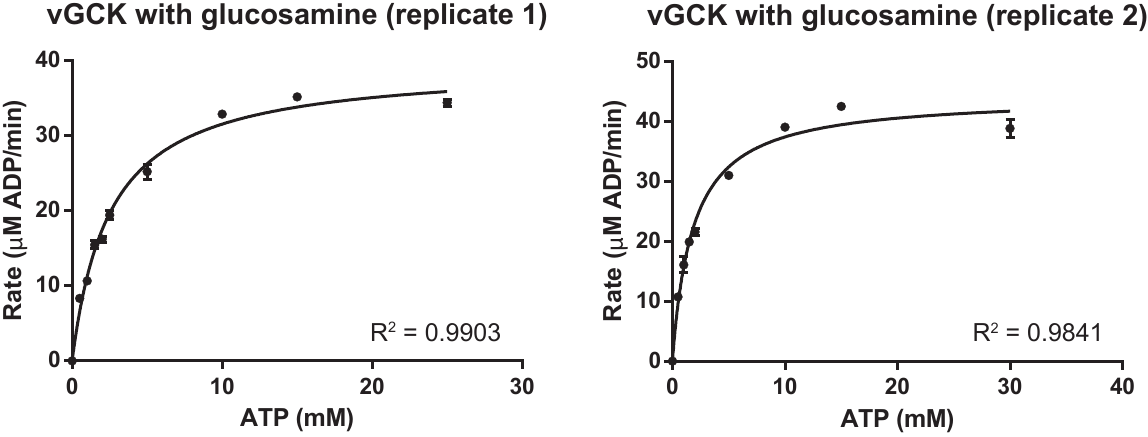

**Figure S18.** Two replicates of steady-state kinetic assays of vGCK with varying ATP concentrations at 50 mM glucosamine. Each data point represents the average ± standard deviation of three technical replicates. Data were fit to the Michaelis-Menten equation.

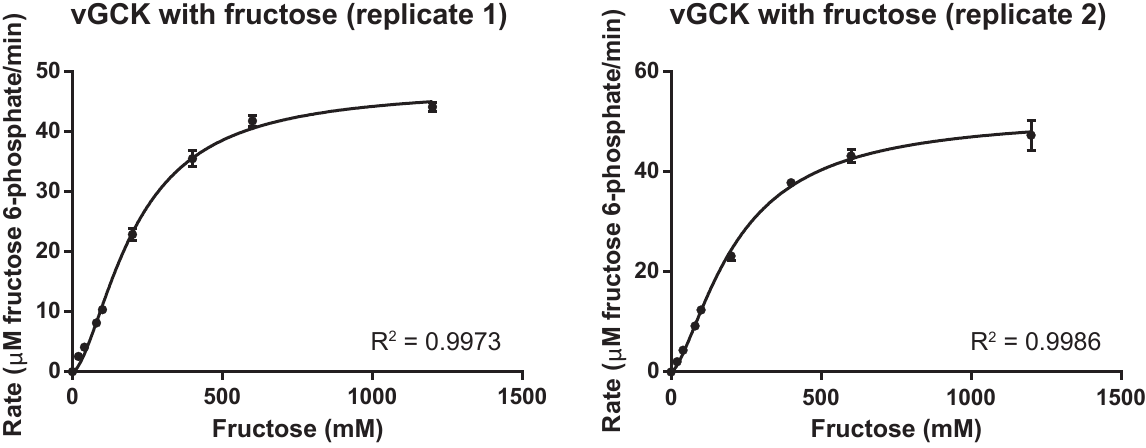

**Figure S19.** Two replicates of steady-state kinetic assays of vGCK with varying fructose concentrations at 10 mM ATP. Each data point represents the average ± standard deviation of three technical replicates. Data were fit to the Hill equation.

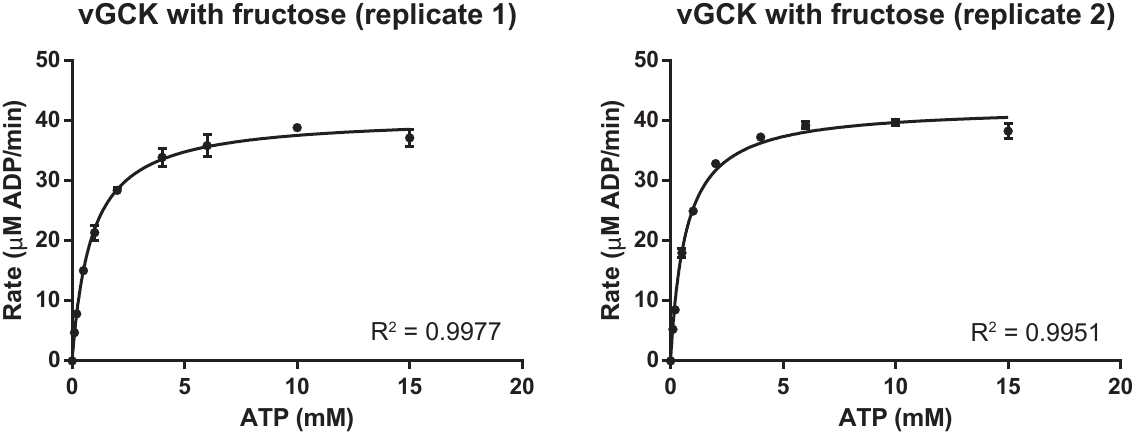

**Figure S20.** Two replicates of steady-state kinetic assays of vGCK with varying ATP concentrations at 800 mM fructose. Each data point represents the average ± standard deviation of three technical replicates. Data were fit to the Michaelis-Menten equation.

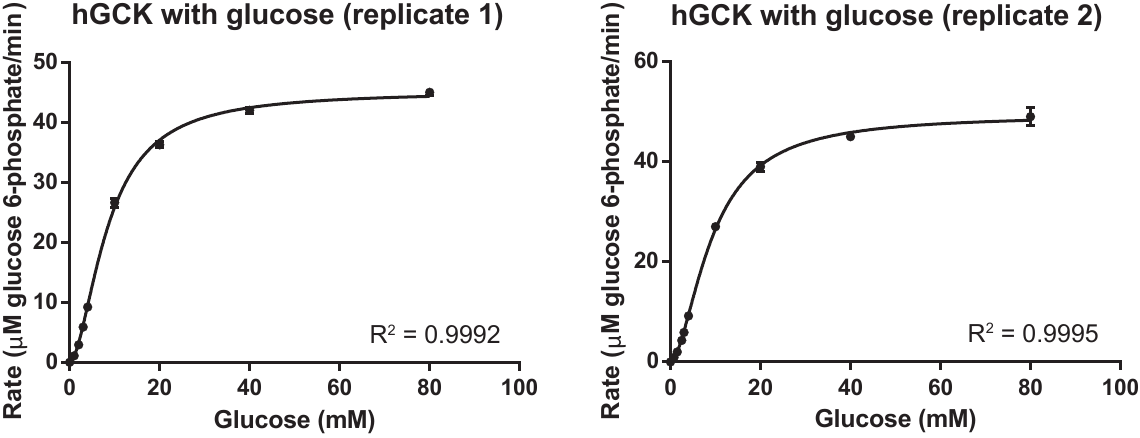

**Figure S21.** Two replicates of steady-state kinetic assays of hGCK with varying glucose concentrations at 10 mM ATP. Each data point represents the average ± standard deviation of three technical replicates. Data were fit to the Hill equation.

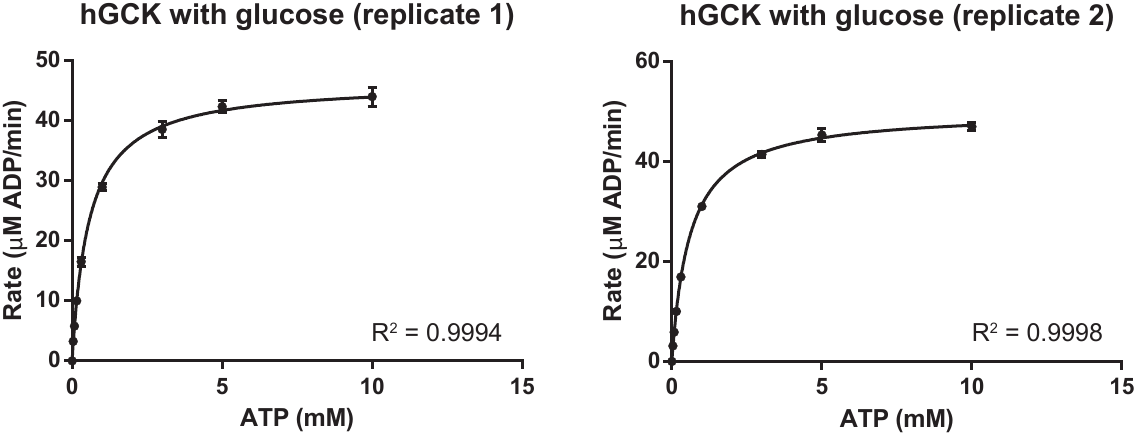

**Figure S22.** Two replicates of steady-state kinetic assays of hGCK with varying ATP concentrations at 100 mM glucose. Each data point represents the average ± standard deviation of three technical replicates. Data were fit to the Michaelis-Menten.

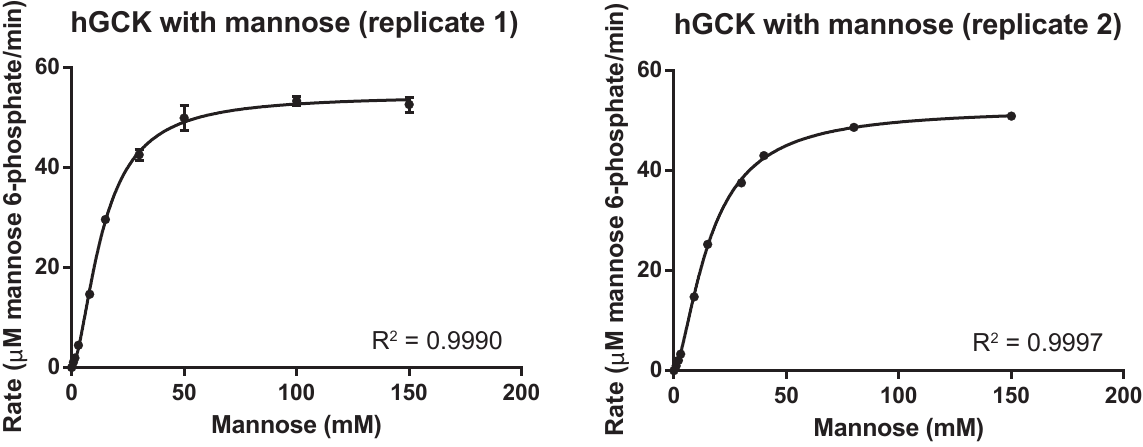

**Figure S23.** Two replicates of steady-state kinetic assays of hGCK with varying mannose concentrations at 10 mM ATP. Each data point represents the average ± standard deviation of three technical replicates. Data were fit to the Hill equation.

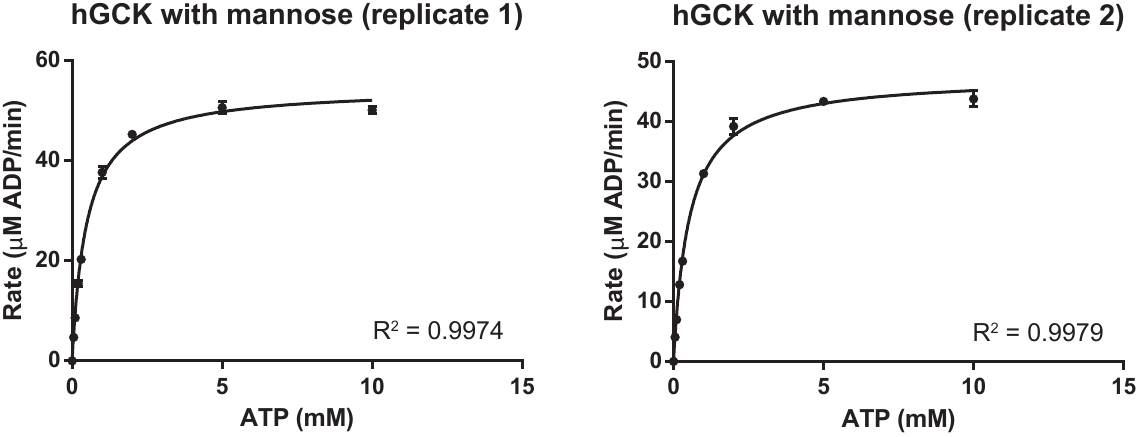

**Figure S24.** Two replicates of steady-state kinetic assays of hGCK with varying ATP concentrations at 150 mM mannose. Each data point represents the average ± standard deviation of three technical replicates. Data were fit to the Michaelis-Menten equation.

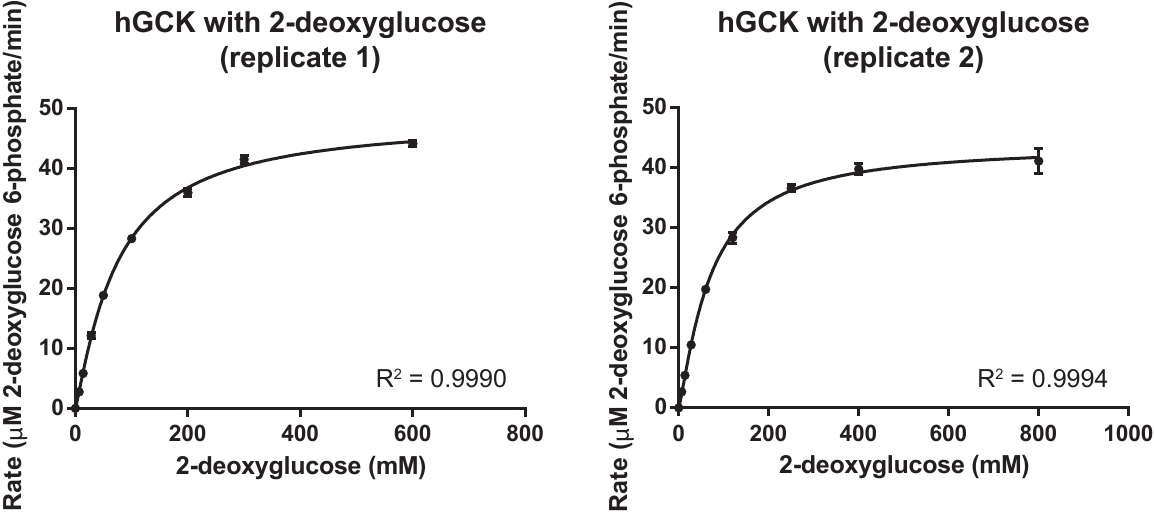

**Figure S25.** Two replicates of steady-state kinetic assays of hGCK with varying 2-deoxyglucose concentrations at 15 mM and 10 mM ATP for replicate 1 and 2, respectively. Each data point represents the average ± standard deviation of three technical replicates. Data were fit to the Hill equation.

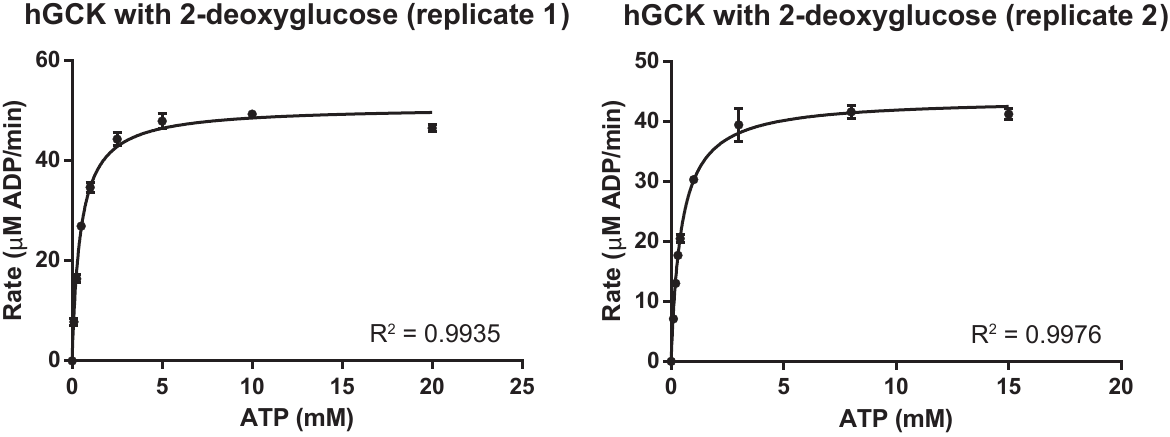

**Figure S26.** Two replicates of steady-state kinetic assays of hGCK with varying ATP concentrations at 700 mM 2-deoxyglucose. Each data point represents the average ± standard deviation of three technical replicates. Data were fit to the Michaelis-Menten equation.

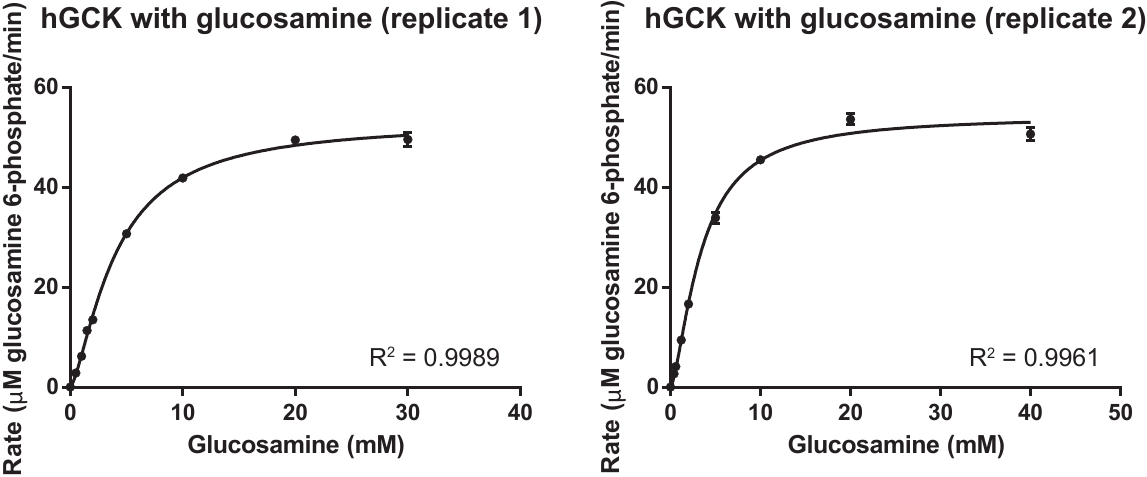

**Figure S27.** Two replicates of steady-state kinetic assays of hGCK with varying glucosamine concentrations at 15 mM and 20 mM ATP for replicate 1 and 2, respectively. Each data point represents the average ± standard deviation of three technical replicates. Data were fit to the Hill equation.

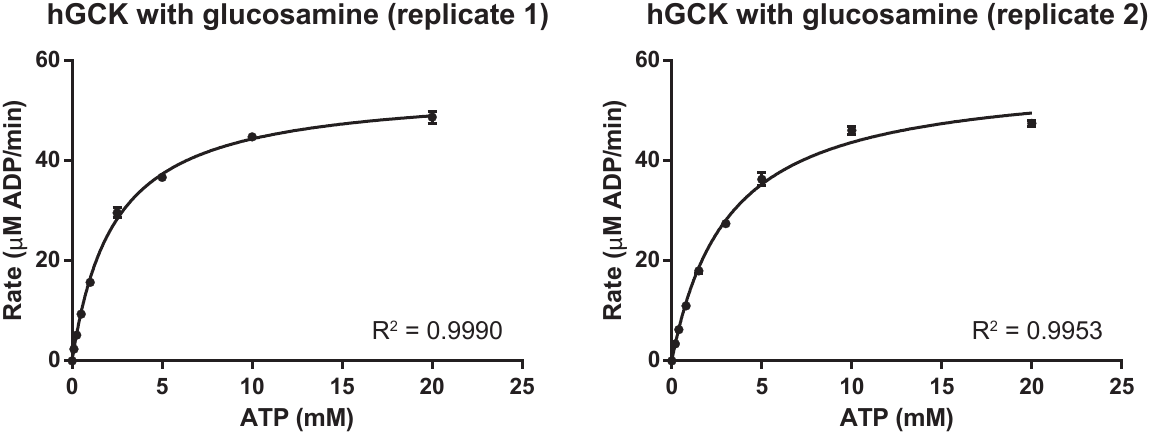

**Figure S28.** Two replicates of steady-state kinetic assays of hGCK with varying ATP concentrations at 30 mM glucosamine. Each data point represents the average ± standard deviation of three technical replicates. Data were fit to the Michaelis-Menten equation.

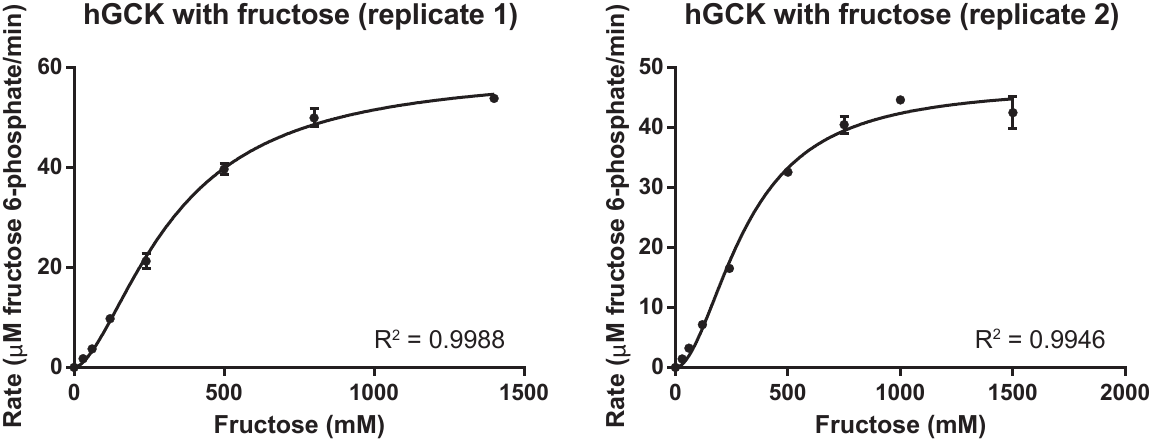

**Figure S29.** Two replicates of steady-state kinetic assays of hGCK with varying fructose concentrations at 10 mM ATP. Each data point represents the average ± standard deviation of three technical replicates. Data were fit to the Hill equation.

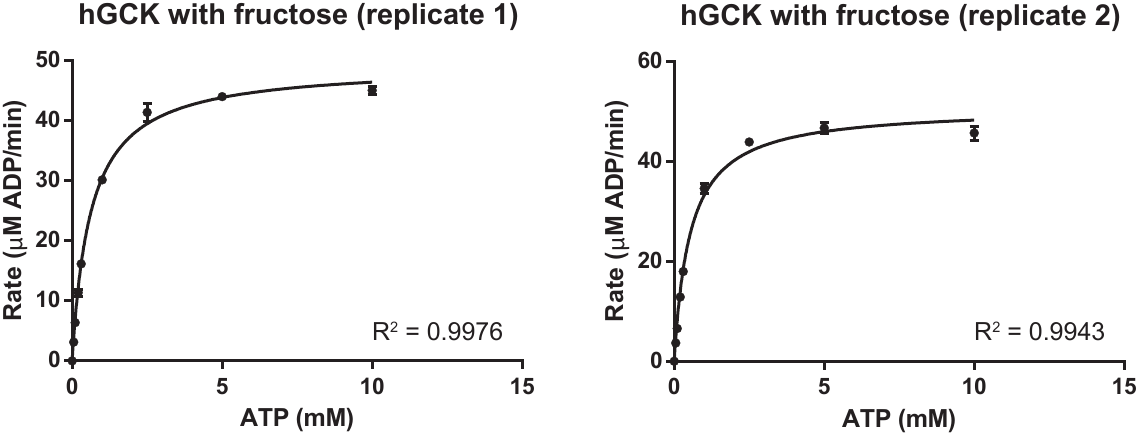

**Figure S30.** Two replicates of steady-state kinetic assays of hGCK with varying ATP concentrations at 1.5 M fructose. Each data point represents the average ± standard deviation of three technical replicates. Data were fit to the Michaelis-Menten equation.

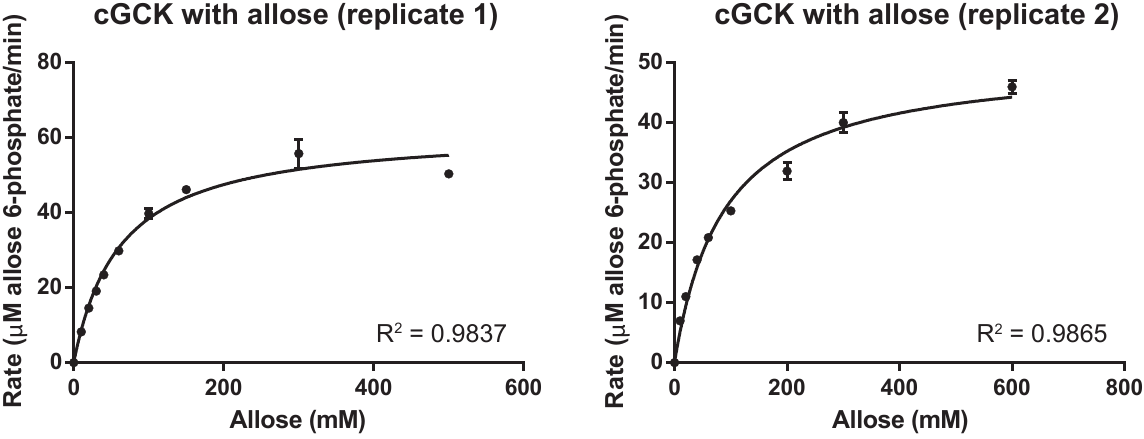

**Figure S31.** Two replicates of steady-state kinetic assays of cGCK with varying allose concentrations at 10 mM ATP. Each data point represents the average ± standard deviation of three technical replicates. Data were fit to the Michaelis-Menten equation.

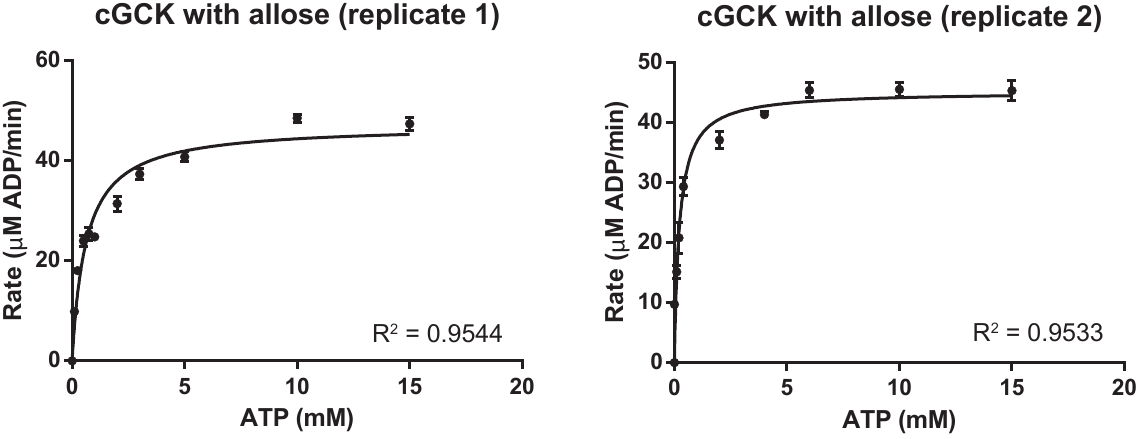

**Figure S32.** Two replicates of steady-state kinetic assays of cGCK with varying ATP concentrations at 400 mM allose. Each data point represents the average ± standard deviation of three technical replicates. Data were fit to the Michaelis-Menten equation.

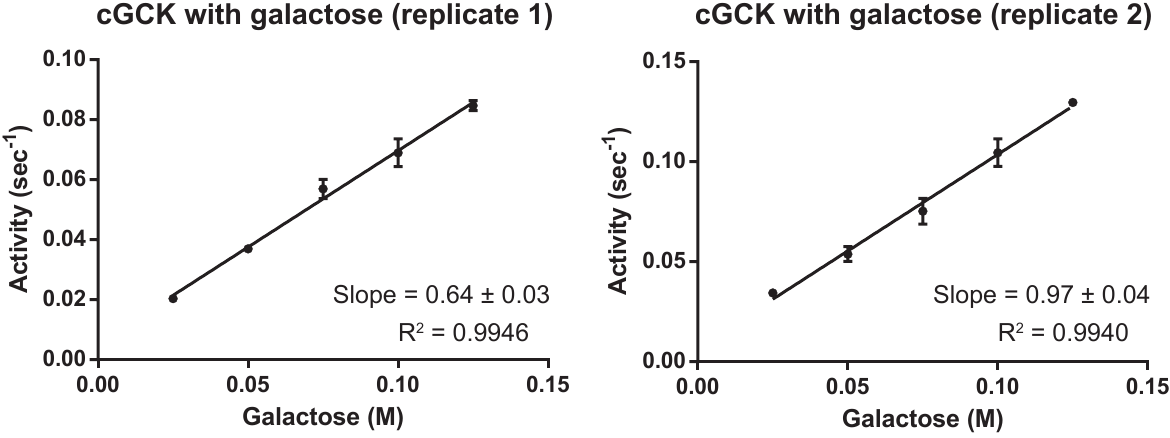

**Figure S33.** Two replicates of steady-state kinetic assays of cGCK with varying galactose concentrations at 15 mM ATP to estimate catalytic efficiency, *k*_cat_/*K*_m_. Each data point represents the average ± standard deviation of three technical replicates. Data were fit to the linear regression equation.

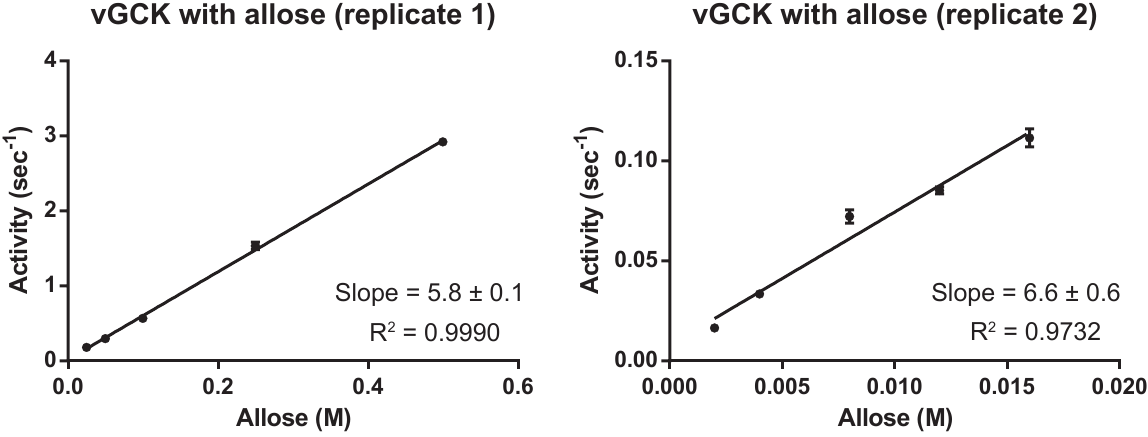

**Figure S34.** Two replicates of steady-state kinetic assays of vGCK with varying allose concentrations at 15 mM ATP to estimate catalytic efficiency, *k*_cat_/*K*_m_. Each data point represents the average ± standard deviation of three technical replicates. Data were fit to the linear regression equation.

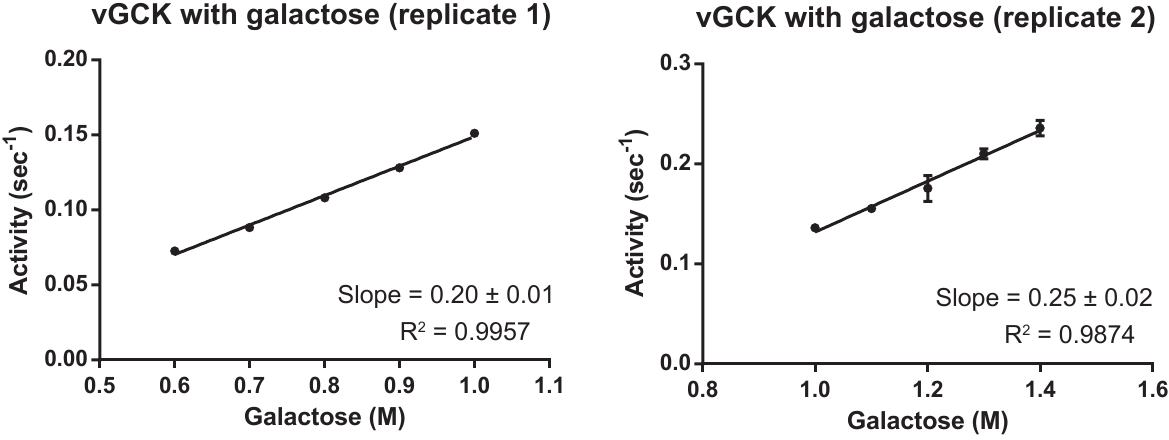

**Figure S35.** Two replicates of steady-state kinetic assays of vGCK with varying galactose concentrations at 15 mM ATP to estimate catalytic efficiency, *k*_cat_/*K*_m_. Each data point represents the average ± standard deviation of three technical replicates. Data were fit to the linear regression equation.

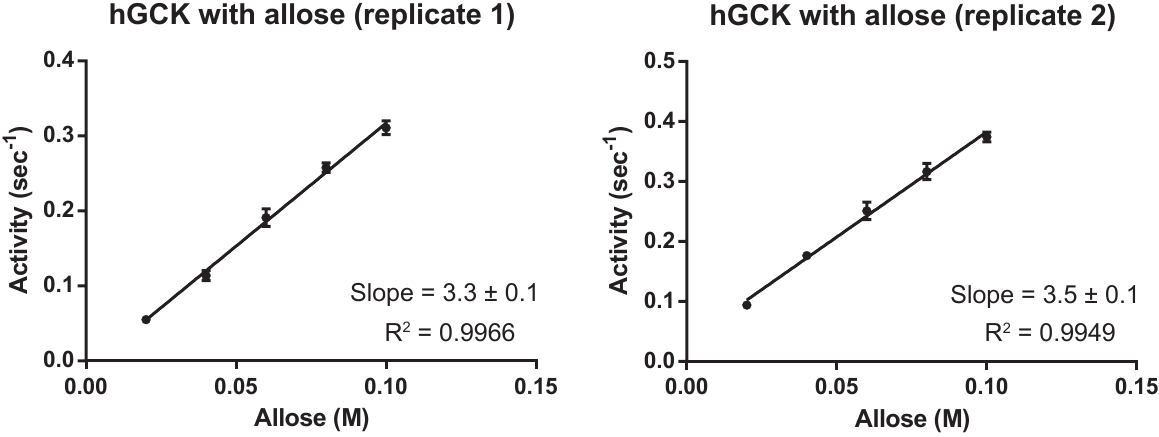

**Figure S36.** Two replicates of steady-state kinetic assays of hGCK with varying allose concentrations at 15 mM ATP to estimate catalytic efficiency, *k*_cat_/*K*_m_. Each data point represents the average ± standard deviation of three technical replicates. Data were fit to the linear regression equation.

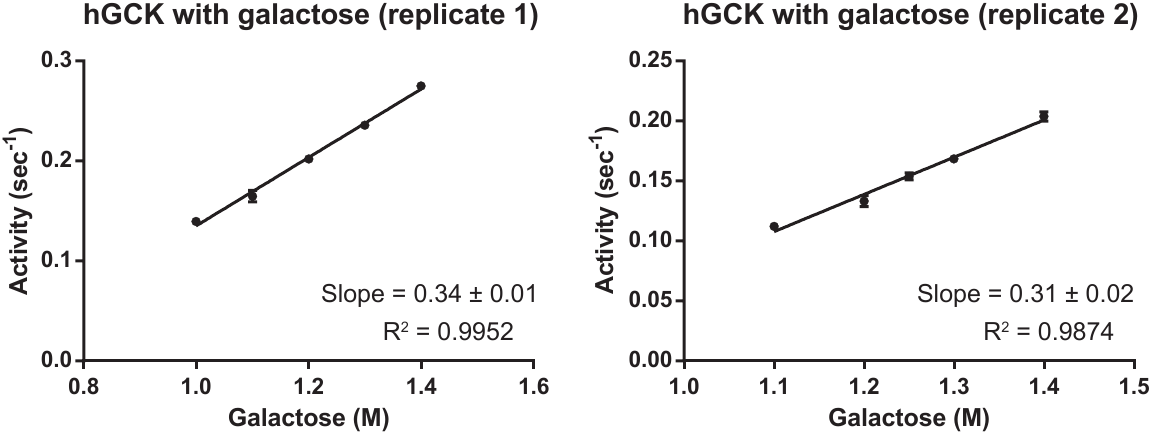

**Figure S37.** Two replicates of steady-state kinetic assays of hGCK with varying galactose concentrations at 15 mM ATP to estimate catalytic efficiency, *k*_cat_/*K*_m_. Each data point represents the average ± standard deviation of three technical replicates. Data were fit to the linear regression equation.

A)

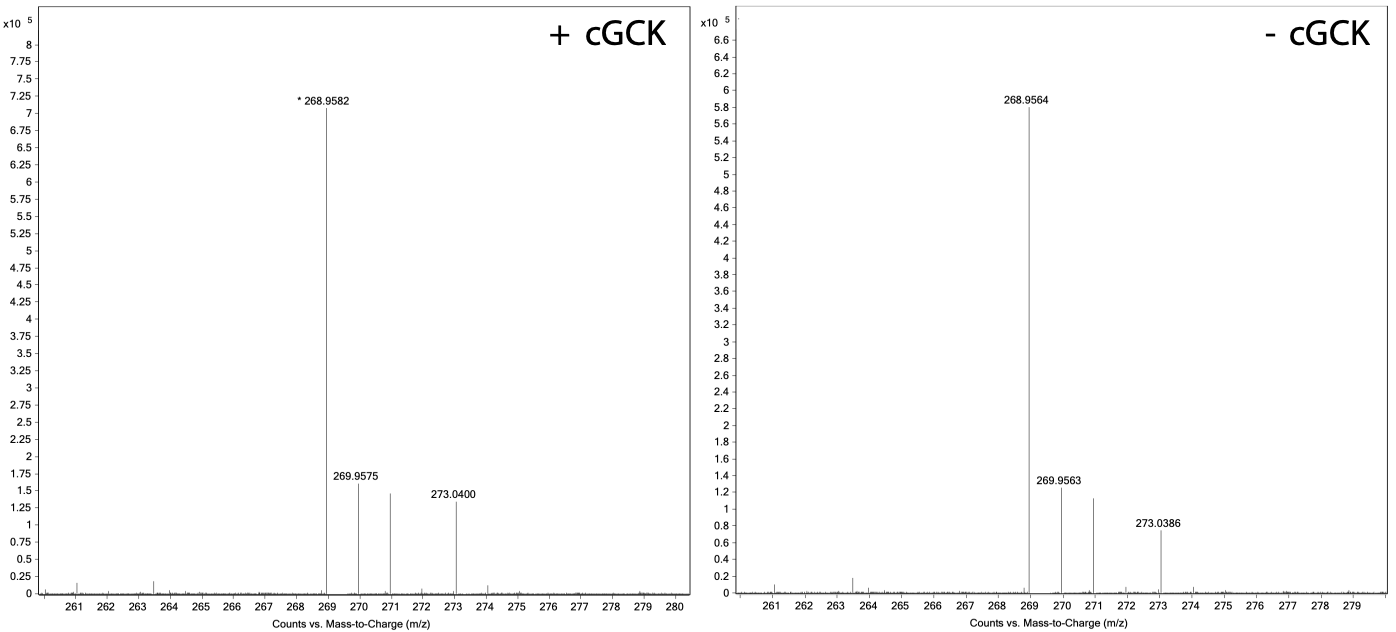

B)

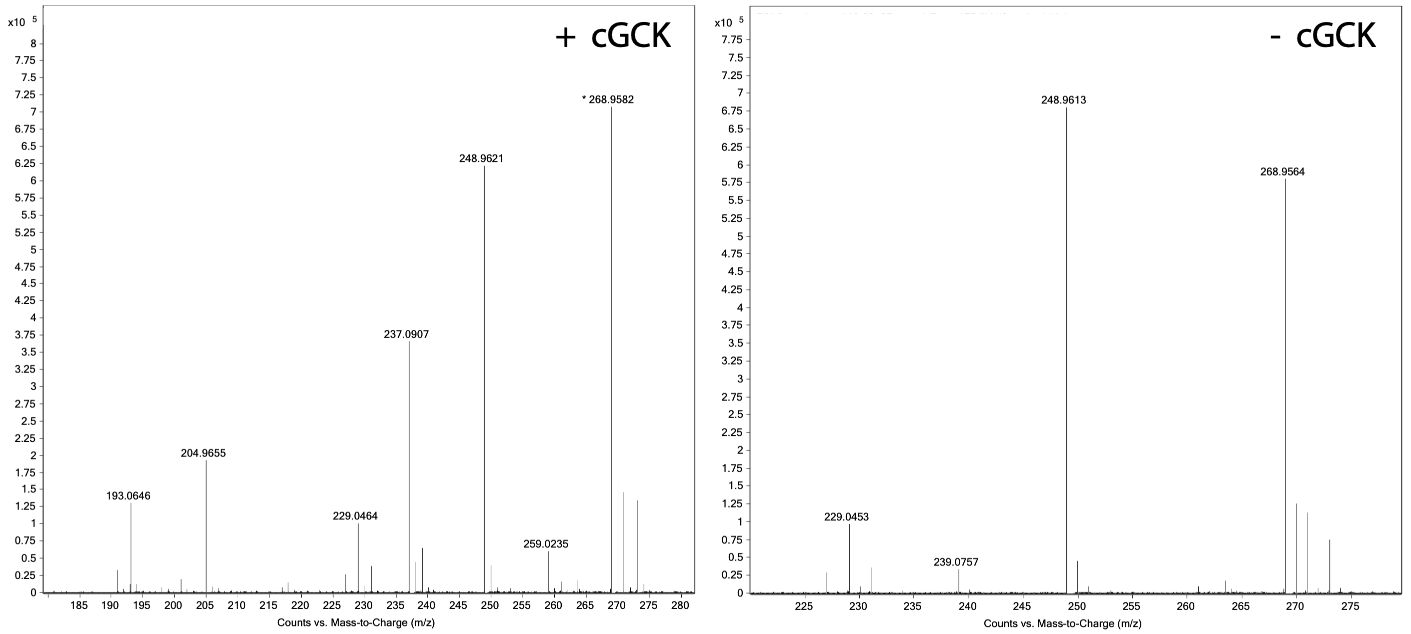

**Figure S38.** Analysis of reaction mixtures containing (left) and lacking cGCK (right) via mass spectrometry to probe the enzymatic production of methyl-glucopyranose 6-phosphate. A) Methyl-glucopyranose is detected at 273 m/z in both samples suggesting that product formation is spontaneous. B) A peak at 259 m/z, correpsponding to the mass to charge ratio of glucose 6-phosphate or analogs thereof, appears in the spectra containing cGCK (left) but is absent in the spectra lacking cGCK (right).

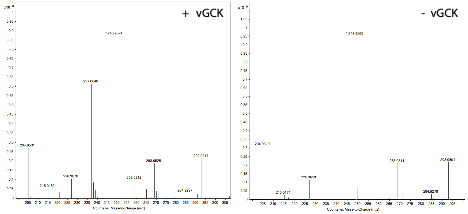

**Figure S39.** Analysis of reaction mixtures containing (left) and lacking vGCK (right) via mass spectrometry to probe the enzymatic production of galactose 6-phosphate. Galactose 6-phosphate (259 m/z) is produced in the presence presence of vGCK but is lacking in the absence of vGCK.

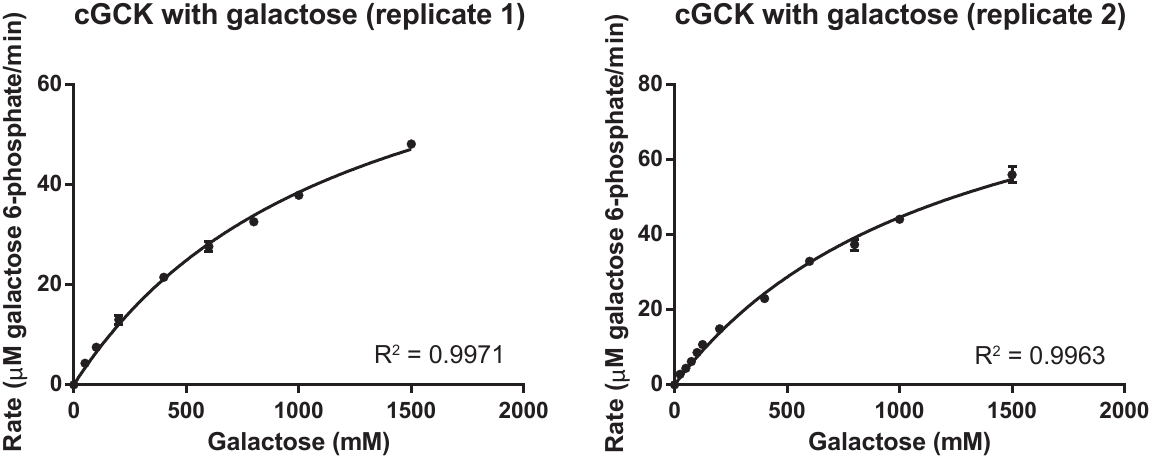

**Figure S40.** Two replicates of steady-state kinetic assays of cGCK with varying galactose concentrations at 15 mM ATP to estimate individual *K*_m_ and *k*_cat_ values. Each data point represents the average ± standard deviation of three technical replicates. Data were fit to the Michaelis-Menten equation.

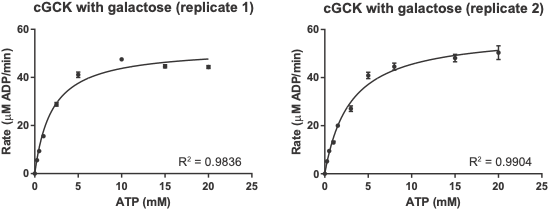

**Figure S41.** Two replicates of steady-state kinetic assays of cGCK with varying ATP concentrations at 1.5 M galactose. Each data point represents the average ± standard deviation of three technical replicates. Data were fit to the Michaelis-Menten equation.
